# Overexpression of mig-6 in cartilage induces an osteoarthritis-like phenotype in mice

**DOI:** 10.1101/764142

**Authors:** Melina Bellini, Michael A. Pest, M. Miranda-Rodrigues, JW Jeong, Frank Beier

## Abstract

**Background:** Osteoarthritis (OA) is the most common form of arthritis and characterized by degeneration of articular cartilage. Mitogen-inducible gene 6 (Mig-6) has been identified as a negative regulator of the Epidermal Growth Factor Receptor (EGFR). Cartilage-specific *Mig-6* knockout (KO) mice display increased EGFR signaling, an anabolic buildup of articular cartilage and formation of chondro-osseous nodules. Since our understanding of the EGFR/Mig-6 network in cartilage remains incomplete, we characterized mice with cartilage-specific overexpression of Mig-6 in this study.

**Methods:** Utilizing knee joints from cartilage-specific *Mig-6* overexpressing (*Mig-*^*6over/over*^) mice (at multiple time points), we evaluated the articular cartilage using histology, immunohistochemical staining and semi-quantitative OARSI scoring at multiple ages. MicroCT analysis was employed to examine skeletal morphometry, body composition, and bone mineral density.

**Results:** Our data show that cartilage-specific *Mig-6* overexpression did not cause any major developmental abnormalities in articular cartilage, although *Mig-6*^*over/over*^ mice have slightly shorter long bones compared to the control group. Moreover, there was no significant difference in bone mineral density and body composition in any of the groups. However, our results indicate that *Mig-6*^*over/over*^ male mice show accelerated cartilage degeneration at 12 and 18 months of age. Immunohistochemistry for SOX9 demonstrated that the number of positively stained cells in *Mig-6*^*over/over*^ mice decreased relative to controls. Immunostaining for MMP13 staining is increased in areas of cartilage degeneration in *Mig-6*^*over/over*^ mice. Moreover, staining for phospho-EGFR (Tyr-1173) and lubricin (PRG4) was decreased in the articular cartilage of *Mig-6*^*over/over*^ mice.

**Conclusion:** Overexpression of *Mig-6* in articular cartilage causes no major developmental phenotype; however these mice develop earlier OA during aging. These data demonstrate that Mig-6/EGFR pathways is critical for joint homeostasis and might present a promising therapeutic target for OA.

## INTRODUCTION

Osteoarthritis (OA), a chronic degenerative joint disease, is the most common form of arthritis. OA affects nearly five million Canadians currently (1), but that number will grow to more than 10 million by 2040 (2). This statistic is alarming, considering the disability, the loss of quality of life, and the costs to the health system generated by OA. Currently, there are pharmacological treatments available to manage OA symptoms such as pain (3–5) as well as surgical joint replacement at the end stage of disease (6,7). Unfortunately however, there is no cure for OA. Progressive understanding of the pathophysiology of OA suggests that the disease is a heterogeneous condition, so further research is needed to direct the clinical approaches to disease management (8).

Recent studies have shown that OA is a multifactorial disease of the whole joint, however its pathogenesis remains still poorly understood (9). Genetic, environmental, and biomechanical factors can accelerate the onset of OA (10). Articular cartilage is a highly specialized tissue that forms the smooth gliding surface of synovial joints, with chondrocytes as the only cellular component of cartilage (11). The homeostasis of the cartilage extracellular matrix (ECM) involves a dynamic equilibrium between anabolic and catabolic pathways controlled by chondrocytes (12). The progression of OA is associated with dramatic alteration in the integrity of the cartilage ECM network formed by a large number of proteoglycans (mostly aggrecan), collagen II, and other non-collagenous matrix proteins (13). In addition, ECM synthesis is regulated by a number of transcriptional regulators involved in chondrogenesis, specifically Sex-determining-region-Y Box 9 (SOX9), L-SOX 5 and SOX6 that regulate type II collagen (*Col2a1*) and Aggrecan (*Acan*) gene expression (14). On the other hand, catabolic events are dominant in OA and cells are exposed to degenerative enzymes such as aggrecanases (e.g. ADAMTS-4, -5) (13,15), collagenases (e.g. MMP-1,-3, -8, -13) (16), and gelatinases (e.g.MMP-2, and MMP-9), all of which have implications in articular cartilage degeneration (17). A number of growth factors (18) play a role in OA pathology, such as transforming growth factor-β (19), BMP-2 (20), Insulin growth factor 1 (IGF-1) (21) fibroblast growth factor (FGF) and others, but the exact regulation of chondrocyte physiology is still not completely understood.

Recent studies in our laboratory (22,23) have identified the epidermal growth factor receptor (EGFR) and its ligand transforming growth factor alpha (TGFα) as possible mediators of cartilage degeneration (24–26). The human *TGFA* gene locus was also strongly linked to hip OA and cartilage thickness in genome-wide association studies (27,28). TGFα stimulates EGFR signaling and activates various cell-signaling pathways in chondrocytes, including extracellular signal-regulated kinase 1 and 2 (ERK1/2) and P13K (phosphoinositide 3-kinase) (29). EGFR signaling plays important roles in endochondral ossification (30,31), growth plate development (30) and cartilage maintenance and homeostasis (32–34), but many aspects of its action in cartilage are still not well understood. However, both protective and catabolic effects of EGFR signaling in OA have been reported, suggesting context-specific roles of this pathway (35).

Mitogen-inducible gene 6 (Mig-6) is also known as Gene 33, ErbB receptor feedback inhibitor 1 (ERRFI1), or RALT, and is found in the cytosol (36). *Mig-6* protein binds to and inhibits EGFR signaling through a two-tiered mechanism: suppression of EGFR catalytic activity and receptor down-regulation (37). Interestingly, various studies have reported that loss of Mig-6 induces the onset of OA-like symptoms in mice (36,38–40). Cartilage-specific (Col2-Cre) knockout of *Mig-6* mice results in formation of chondro-osseous nodules in the knee, but also increased thickness of articular cartilage in the knee, ankle, and elbow (41). Prx1-cre-mediated knockout of *Mig-6* results in a similar phenotype as that observed in cartilage-specific knockout mice(42). These phenotypes appeared to be caused by an increase in chondrocyte proliferation in articular cartilage, supported by increased expression of Sox9 and EGFR activation in cartilage (42). Since our studies suggest dosage- and/or context-specific roles of EGFR signaling in the process of cartilage degeneration in OA, in this study we used a Col2a1 promoter–driven Cre/lox system to examine effects of Mig-6 overexpression specifically in articular cartilage.

## Materials and Methods

### Generation of Mig-6 overexpression mice

*Mig-6* overexpression animals on a mixed C57Bl/6 and agouti mouse background, with the overexpression cassette in the Rosa26 locus (43) and bred for 10 generations into a C57Bl/6 background. Transcription of *Mig-6* is under the control of a ubiquitously expressed chicken beta actin-cytomegalovirus hybrid (CAGGS) promoter, but blocked by a “Stop Cassette” flanked by LoxP sites (LSL) (43). *Mig-6* overexpression mice were bred to mice carrying the Cre recombinase gene under the control of the Collagen 2 promoter (44), to induce recombination and removal of the Stop Cassette specifically in cartilage. Throughout the manuscript, animals for homozygote overexpression of Mig-6 from both alleles are termed *Mig-6* ^*over/over*^ *(Mig-6* ^*over/over*^*Col2a1-Cre*^*+/-*^), while control mice are identical but without the Cre gene (noted as “control” in this manuscript for simplicity). Mice were group housed (at least 1 pair of littermate matched control and overexpression animals), on a standard 12 hour light/dark cycle, without access to running wheels, and with free access to mouse chow and water. Animals were weighed prior to euthanization by asphyxiation with CO2. All animal experiments were done in accordance with the Animal Use Subcommittee at the University of Western Ontario and conducted in accordance with guidelines from the Canadian Council on Animal Care.

### Genotyping

Genotype was determined by polymerase chain reaction (PCR) analysis using DNA processed from biopsy samples of ear tissue from mice surviving to at least 21 days of age. PCR strategy: Primer set P1 and P2 can amplify a 300 bp fragment from the wild-type allele, whereas P1 and P3 can amplify a 450 bp fragment from the targeted ROSA26 locus allele (43) (Supp. Figure/Table 1).

### Histopathology of the knee

Limbs from *Mig-6* ^*over/over*^ and control mice were harvested and fixed in 4% paraformaldehyde (Sigma) for 24 hours and decalcified in ethylenediaminetetraacetic acid (5% EDTA in phosphate buffered saline (PBS), pH 7.0. Joints were processed and embedded in paraffin in sagittal or frontal orientation, with serial sections taken at a thickness of 5 μm. Sections were stained with Toluidine Blue (0.04% toluidine blue in 0.2M acetate buffer, pH 4.0, for 10 minutes) for glycosaminoglycan content and general evaluation of articular cartilage. All images were taken with a Leica DFC295 digital camera and a Leica DM1000 microscope.

### Thickness of proximal tibia growth plate

For early developmental time points such as newborn (P0), sagittal knee sections stained with toluidine blue were used to measure the width of the zones of the epiphyseal growth plate in the proximal tibia. The average thickness of the resting and proliferative zones combined was evaluated by taking three separate measurements at approximately equal intervals across the width of the growth plate. The average hypertrophic zone thickness was also measured using 3 different measurements across the width of the growth plate, starting each measurement at the border of the proliferative and hypertrophic zones and ending at the subchondral bone interface. A third average measurement was then taken of the thickness of the entire growth plate. ImageJ Software (v.1.51) (45) was used for all measures, with the observer blinded to the genotype.

### Articular cartilage evaluation

Articular cartilage thickness was measured from toluidine blue-stained frontal sections by a blinded observer. Articular cartilage thickness was measured separately for the non-calcified articular cartilage (measured from the superficial tangential zone to the tidemark) and the calcified articular cartilage (measured from the subchondral bone to the tidemark) across three evenly spaced points from all four quadrant of the joint (medial/lateral tibia and femur) in 4 sections spanning at least 500 μm. ImageJ Software (v.1.51) (45) was used to measure the thickness of articular cartilage.

### Micro-Computerized Tomography (μCT)

Whole body scans were collected in 6 week-, 11 week-, 12 month- and 18 month-old control and *Mig-6* ^*over/over*^ male and female mice. Mice were euthanized and imaged using General Electric (GE) SpeCZT microCT machine (46) at a resolution of 50μm/voxel or 100μm/voxel. GE Healthcare MicroView software (v2.2) was used to generate 2D maximum intensity projection and 3D isosurface images to evaluate skeletal morphology. MicroView was used to create a line measurement tool in order to calculate the bone lengths, femurs lengths were calculated from the proximal point of the greater trochanter to the base of the lateral femoral condyle. Tibiae lengths were measured from the midpoint medial plateau to the medial malleolus. Humerus lengths were measured from the midpoint of the greater tubercle to the center of the olecranon fossa.

### Body Composition Analysis

MicroView software (GE Healthcare Biosciences) was used to analyse the microCT scans at the resolution of 100um/voxel. Briefly, the region of interest (ROI) was used to calculate the mean of air, water and an epoxy-based, cortical bone-mimicking calibrator (SB3; Gammex, Middleton, WI, USA) (1100mg/cm^3^) (47). A different set of global thresholds was applied to measure adipose, lean and skeletal mass (−275, -40 and 280 Hounsfield Units (HU), respectively). Moreover, bone mineral density (BMD) was acquired as the ratio of the average HU (from the value of skeletal region of interest) in order to calculate HU value of the SB3 calibrator, multiplied by the known density of the SB3 as described (46).

### OARSI histopathology scoring

Serial sections through the entire knee joint were scored according to the OARSI histopathology scoring system (48) by two blinded observers on the four quadrants of the knee: lateral femoral condyle (LFC), lateral tibial plateau (LTP), medial femoral condyle (MFC), and medial tibial plateau (MTP). Histologic scoring from 0-6 represent the OA severity, from 0 (healthy cartilage) to 6 (erosion of more than 75% of articular cartilage). Individual scores are averaged across observers and OA severity is shown as described for each graph. Scores were compared between male and female *Mig-6* ^*over/over*^ and control mice at both 12 and 18 months of age. All images were taken with a Leica DFC295 digital camera and a Leica DM1000 microscope.

### Immunohistochemistry

Frontal paraffin sections of knees were used to for immunohistochemical analysis, with slides with ‘no primary antibody’ as control. All sections were deparaffinized and rehydrated as previously described (41,49). Subsequently, the sections were incubated in 3% H2O2 in methanol for 15 minutes to inhibit endogenous peroxidase activity. After rising with water, 5% goat or donkey serum in PBS was applied to reduce nonspecific background staining. Sections were incubated overnight at 4°C with primary antibodies against SOX9 (R&D Systems, AF3075), MMP13 (Protein Tech, Chicago, IL, USA, 18165-1-AP), lubricin (Abcam, ab28484) and phospho-EGFR (phosphoTyr-1173; Cell Signaling Technology). After washing, sections were incubated with horseradish peroxidase (HRP)-conjugated donkey anti-goat or goat anti-rabbit secondary antibody (R&D system and Santa Cruz), before incubation with diaminobenzidine substrate as a chromogen (Dako, Canada). Finally, sections were counterstained with 0.5% methyl green (Sigma) and mounted. Cell density of articular cartilage chondrocytes from 6 and 11 weeks-old male mice was determined by counting all lacunae with evidence of nuclear staining in the lateral and medial femur/tibia using a centered region of interest measuring 200 μm wide and 70 μm deep from the articular surface by a blinded observer. For newborn (P0) animals the region of interest measured 200 μm wide and 100 μm deep from the proliferative zone.

### Statistical Analysis

All statistical analyses were performed using GraphPad Prism (v6.0). Differences between two groups were evaluated using Student’s *t-*test, and Two-Way ANOVA was used to compare 4 groups followed by a Bonferroni multiple comparisons test. All *n* values represent the number of mice used in each group/genotyping.

## RESULTS

### Overexpression of Mig-6 has minor effects on skeletal phenotypes during development

We bred mice for conditional overexpression of Mig-6 (43) to mice expressing Cre recombinase under control of the collagen II promoter. Homozygote mice overexpressed Mig-6 in all collagen II-producing cells (and their progeny) from both Rosa26 alleles and are referred to as *Mig-6* ^*over/over*^ from here on. Control mice do not express Cre. Genomic DNA was extracted from ear notches to identify homozygous mice *Mig-6*^*over/over*^ using standard PCR analysis. Overexpressing mice were obtained at the expected Mendelian ratios (data not shown). Male mutant gained weight at the same rate as controls over the examined at 10 week period, while female *Mig-6*^*over/over*^ mice were slightly lighter than controls starting at 8 weeks of age (Fig. 1A,B). These differences persisted at 12 months of age for female mice, while at 18 months both male and female mutant mice were lighter than their controls (Fig. 1C,D).

**Figure 1).**
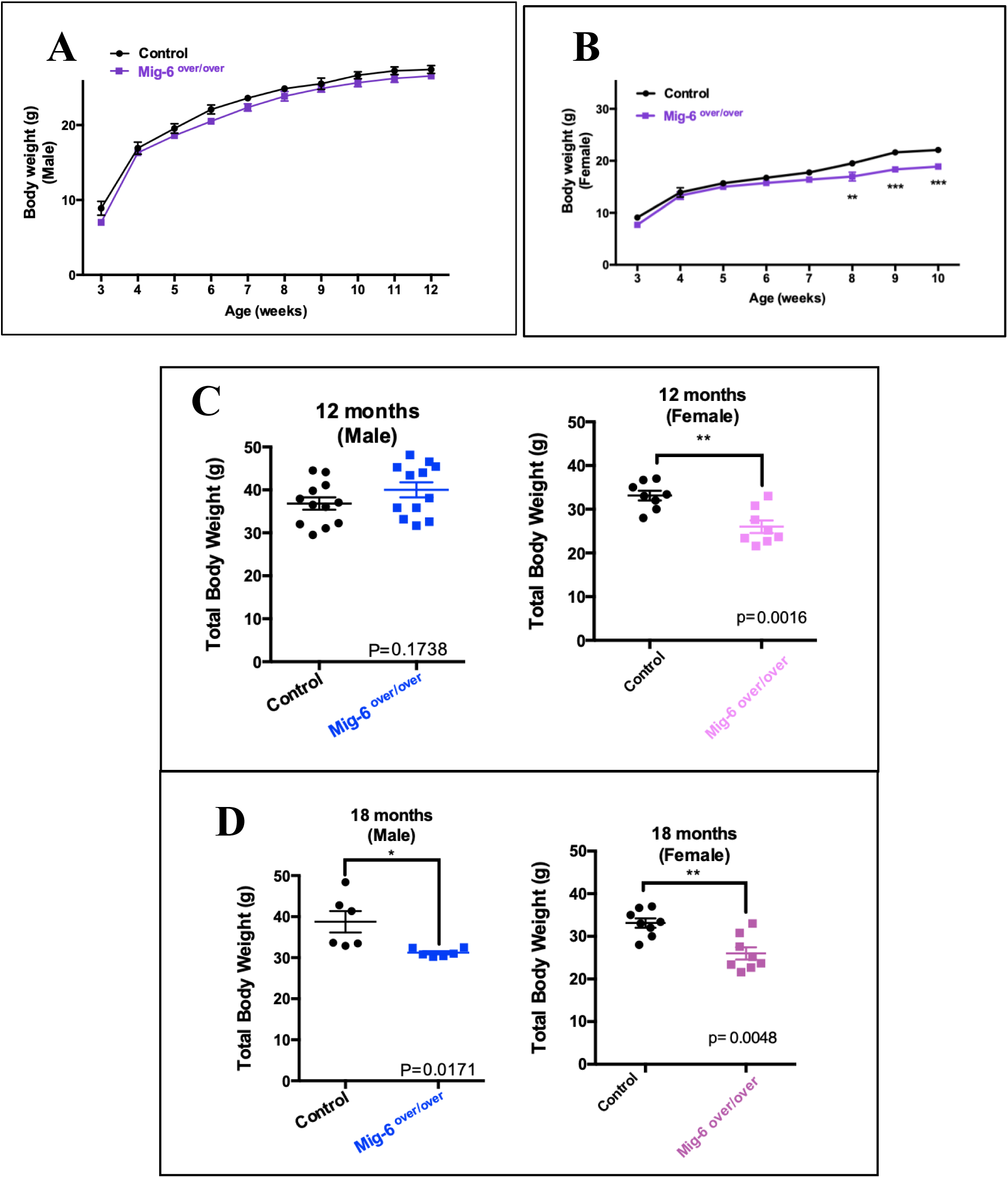
Body weight of control and *Mig-6*^*over/over*^ male and female mice during growth. Body weight of male Mig-6 overexpression mice did not show any significant differences compared to control **(A)** Female Mig-6 overexpression mice showed statistically significant differences compared to control at 8w, 9w and 10w **(B).** Two-Way ANOVA was used with Bonferroni post hoc analysis (n=5/genotyping). Data are presented with mean and error ± SEM (P<0.05). Weights of 12 months old **(C)** and 18 months old **(D)** male and female cartilage specific *Mig-6*^*over/over*^ mice and controls taken immediately prior to sacrifice. Individual data points presented with mean ± SEM (P<0.05). Data analyzed by two tailed student t-tests from 6-12 mice per group (age/genotyping).

Growth plates of post-natal day 0 (P0) *Mig-6* ^*over/over*^ and control mice were analyzed by histology. No major differences in tibia growth plate architecture were seen between genotypes (Fig. 2A). While the length of the total growth plate was slightly reduced in *Mig-6* ^*over/over*^ mice, differences in lengths of either the combined resting/proliferative or hypertrophic zones were not statistically significant (Fig. 2B-D).

**Figure 2).**
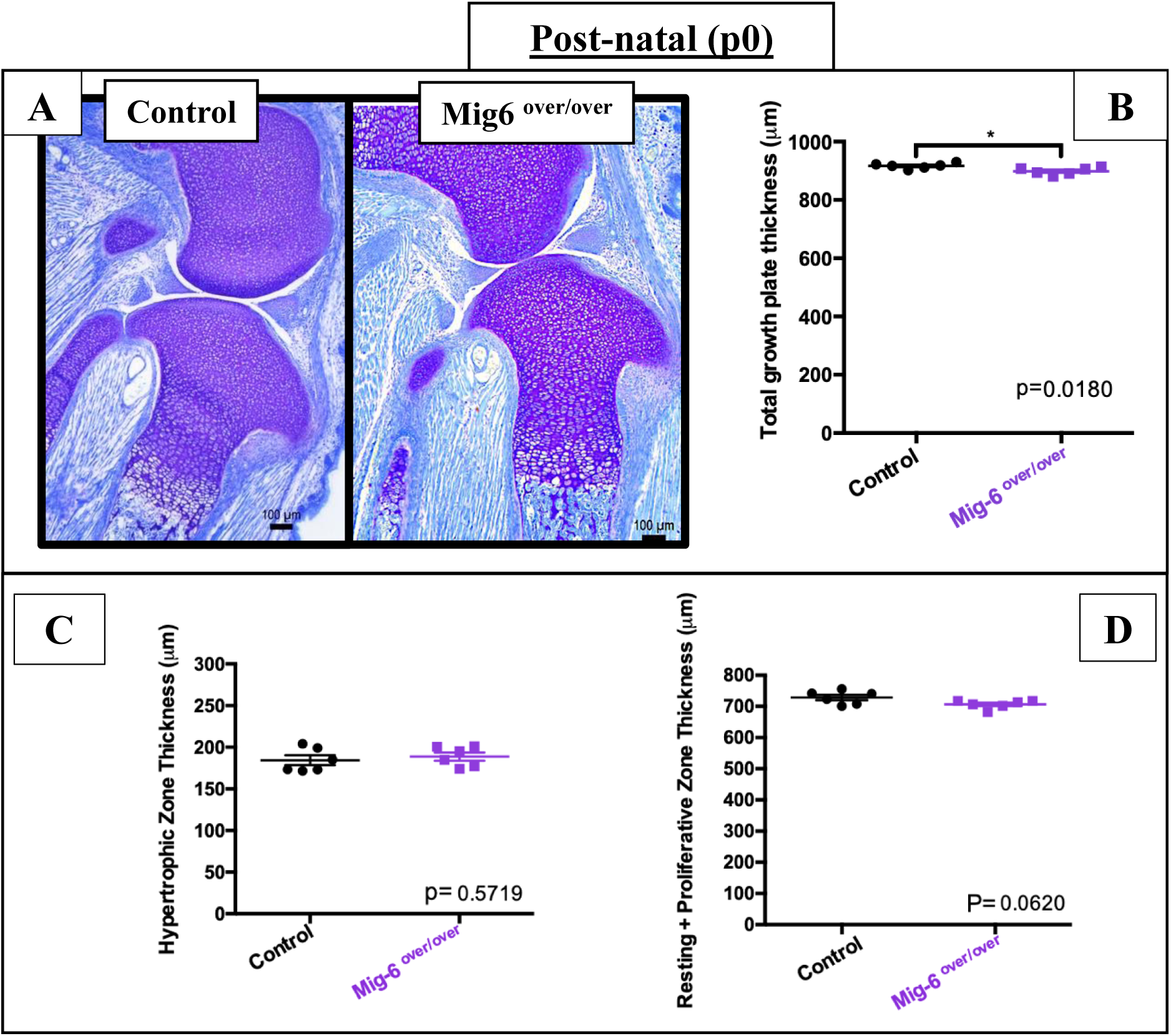
Cartilage-specific Mig6-overexpressing mice display no major developmental phenotype. Representative toluidine Blue staining on postnatal day 0 (P0) of *Mig-6*^*over/over*^ **(A)** and control animals. Thickness of total proximal tibia growth plates in the mice containing articular cartilage specific mitogen inducible gene 6 overexpression (n=6), when compared to age matched controls (n=6) was significantly decreased when analyzed by two tailed student t-tests **(B)**; The average of hypertrophic zone thickness from postnatal day 0 in the mice containing articular cartilage specific mitogen inducible gene 6 overexpression had mean of 188.9 µm and control mice had mean of 184.4 µm **(C).** The thickness of the combined resting and proliferative zones from the control had mean of 728 µm and *Mig-6*^*over/over*^ 706.5 µm *Mig-6*^*over/over*^**(D).** Therefore, there was no significant differences within the groups. Individual data points presented with mean ± SEM analyzed by two tailed student t-tests; (P<0.05).

### Mice overexpressing Mig-6 have shorter long bones than control mice

Skeletal morphology and bone length were examined by microCT mice at the ages of 6 and weeks, and 12 and 18 months. Scans of *Mig-6* ^*over/over*^ male and female mice and their controls were used to generate 3D isosurface reconstructions of 100μm/voxel uCT scans, in order to measure long bones lengths (femurs, humeri, and tibiae) in GE MicroView v2.2 software. Mutant bones were slightly shorter throughout life, with the exception of the male humeri at 12 months that did not show any statistically significant difference (Fig. 3). In contrast, male mice did not show any differences in bone mineral density at 11 weeks, 12 months, or 18 months, compared to controls (Suppl. Fig. 2). In addition, no differences in body mass composition were seen in male mutant and control mice at 11 weeks, 12 months, and 18 months of age (Suppl. Fig. 3).

**Figure 3).**
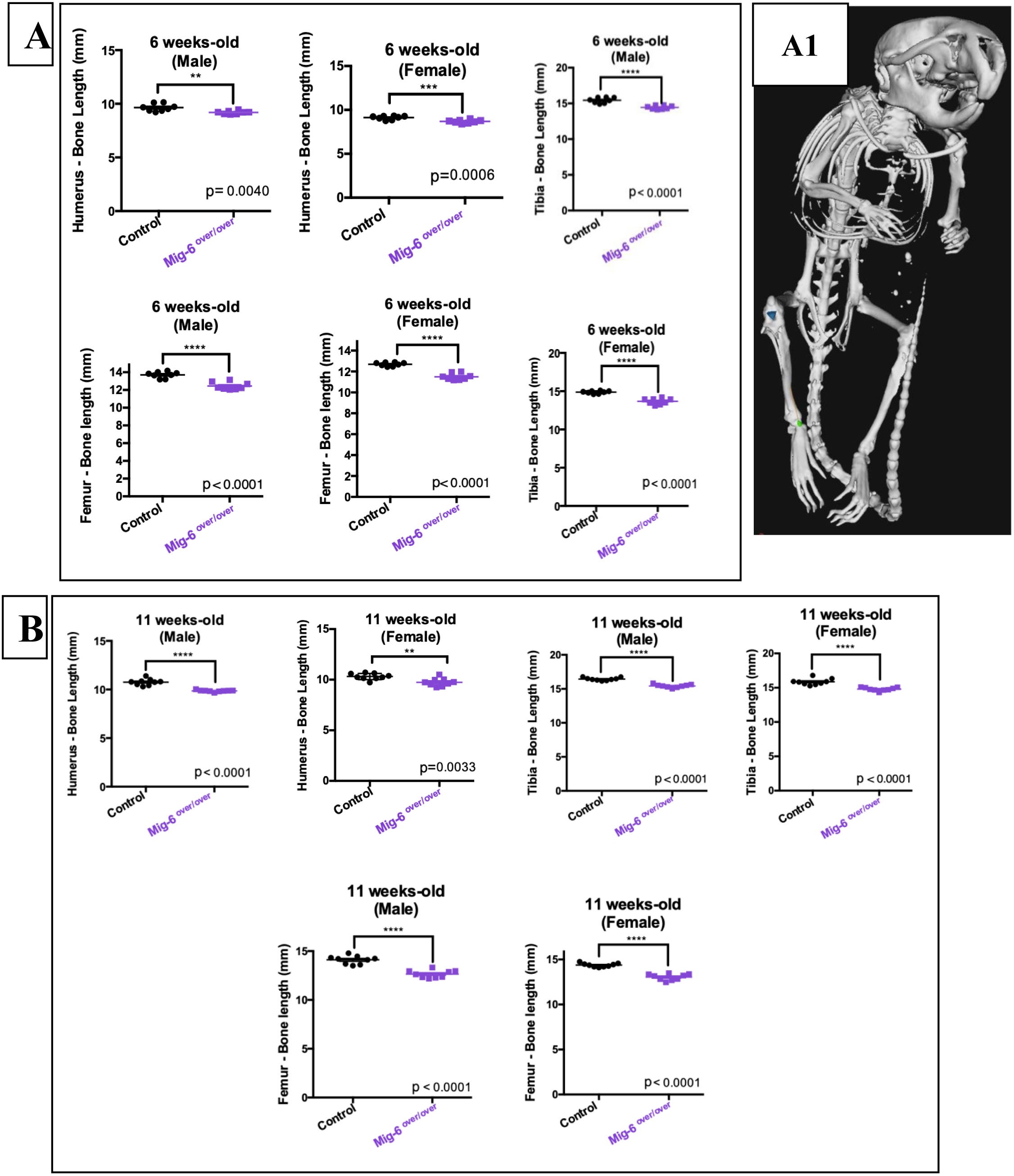
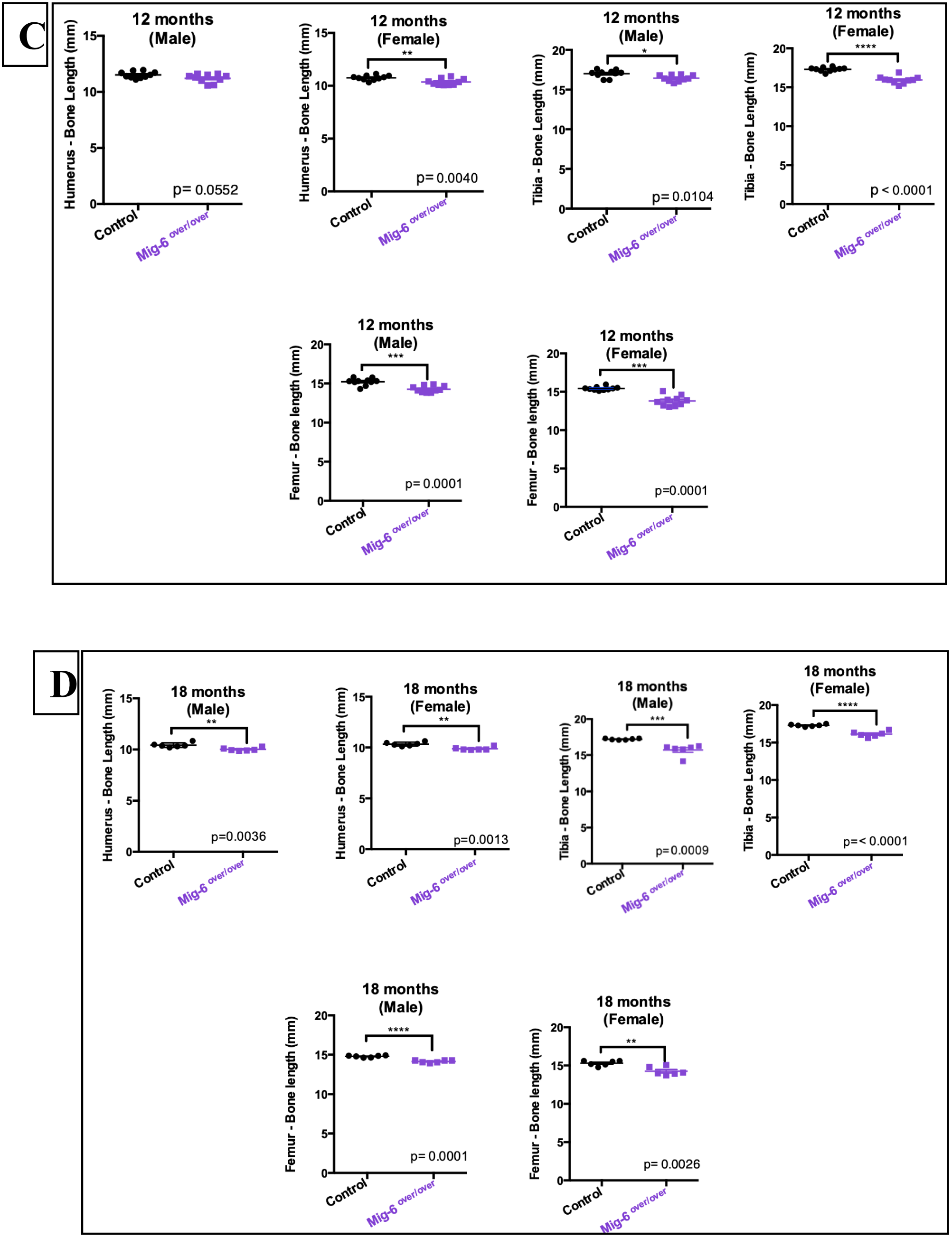
Long bone lengths of Mig-6 overexpression are significantly shorter than control long bone lengths during growth and aging. The lengths of right humeri, tibiae and femora were measured on microCT scan of mice in each different time-points of age using GE MicroView software. **(A)** 6 weeks-old male and female control and Mig-6 overexpressors. **(B)** 11 weeks-old male and female control and Mig-6 overexpressors. **(C)** 12 months-old male and female control and Mig-6 overexpressors. **(D)** 18 months-old male and female control and Mig-6 overexpressors. **(A1)** Representative 3D isosurface reconstructions of 100μm/voxel µCT scans. There were statistically significant differences among control and *Mig-6*^*over/over*^ male and female groups. Individual data points presented with mean ± SEM (P<0.05). Data analyzed by two tailed student t-tests from 6-12 mice per group (age/gender).

### Mig-6 overexpressing mice have healthy articular cartilage during skeletal maturity

We next examined articular cartilage morphology in 11 week-old mutant and control mice using toluidine blue stained paraffin frontal knee sections (Fig. 4A-B). The average thickness of the calcified articular cartilage and non-calcified articular cartilage in the lateral femoral condyle (LFC), lateral tibial plateau (LTP), medial femoral condyle (MFC), and medial tibial plateau (MTP) from control and *Mig-6* ^*over/over*^ male (Fig. 4C-D) and female (Suppl. Fig. 4A-D) mice did not show statistically significant differences. Histological analyses of knee sections from male and female mice did not show any loss of proteoglycan, fibrillation or erosion in the articular cartilage of mutant mice.

**Figure 4).**
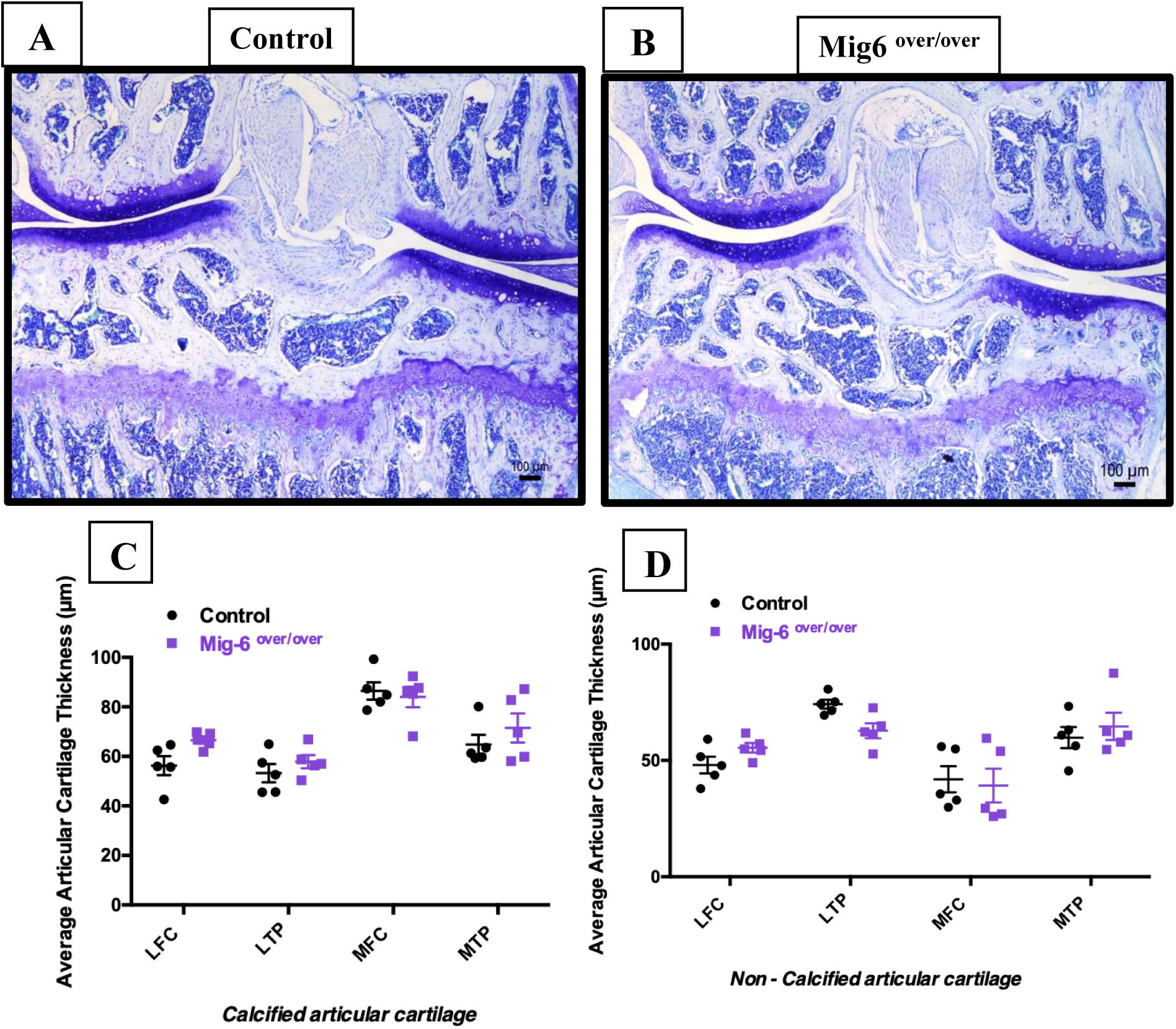
Articular cartilage from 11 weeks-old *Mig-6*^*over/over*^ male mice appeared healthy during skeletal maturity. Representative (n=5/group, toluidine blue) stained frontal sections of knee joints from 11-week-old control **(A)** and Mig-6over **(B).** Mig-6 overexpressors mice show similar articular cartilage thickness when compared to controls at 11 weeks-old male mice. The average thickness of the calcified articular cartilage **(C)** and non-calcified articular cartilage **(D)** in the lateral femoral condyle (LFC), lateral tibial plateau (LTP), medial femoral condyle (MFC), medial tibial plateau (MTP) was measured. Individual data points presented with mean ± SEM. Data analyzed by two-way ANOVA (95% CI) with Bonferroni post-hoc test. Scale bar = 100µm.

### Overexpression of Mig-6 in cartilage induces an osteoarthritis-like phenotype in mice during aging

Since aging is a primary risk factor in OA (50), we next examined knee joints in 12 and 18 month-old control and *Mig-6* ^*over/over*^ mice. Toluidine blue stained sections were evaluated by two blinded observers, using OARSI recommendations (48). At 12 month of age, male control mice showed minor signs of cartilage damage, such as loss of proteoglycan staining, but no significant structural degeneration (Fig. 5A). However, seven of nine *Mig-6* ^*over/over*^ male mice showed more extensive cartilage damage in their medial side (erosion to the calcified layer lesion for 25% to 50% of the medial quadrant). OARSI scoring confirmed increased OA-like damage in mutant mice (Fig. 5C). Similarly, at 18 months of age the male control group showed minimal cartilage degeneration in 3 of 6 mice (Fig. 6A). *Mig-6* ^*over/over*^ male mice showed more severe cartilage erosion in the medial tibial plateau in 4/6 animals. This result was again supported by significantly increased OARSI cartilage damage scores (Fig. 6C). Moreover, for the female group at 12 months, control mice did not show cartilage damage in any quadrant of the knee. *Mig-6* ^*over/over*^ female mice showed sign of OA-like cartilage damage in 3/8 animals (Supplementary Fig. 5). In addition, at 18 months of age, female control mice showed healthy cartilage, and 4/8 *Mig-6* ^*over/over*^ female mice showed some proteoglycan loss and cartilage degeneration on the medial side (Supplementary Fig. 6).

**Figure 5).**
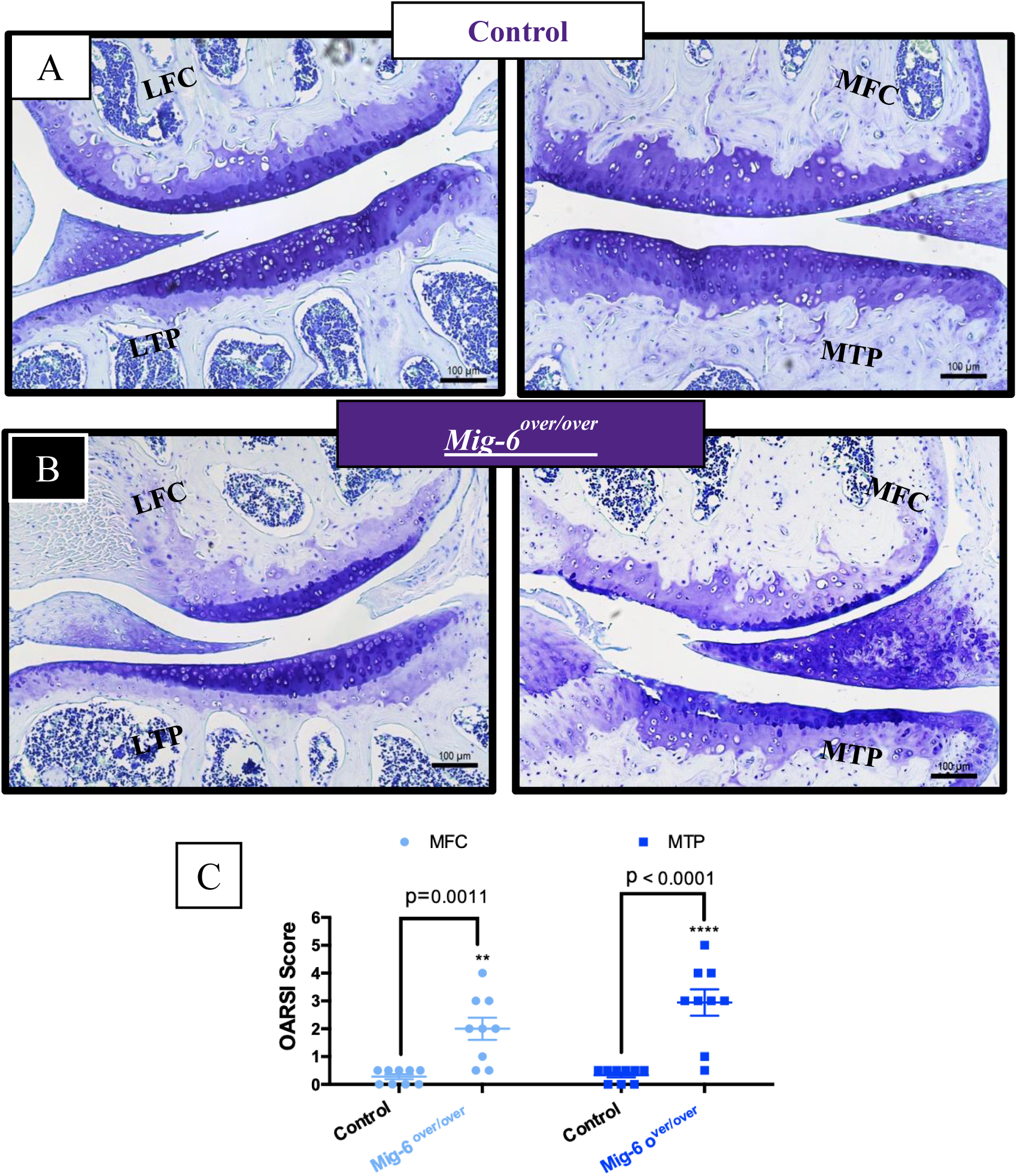
12 months old *Mig-6*^*over/over*^ male mice develop OA-like cartilage degeneration. Representative images of Toluidine Blue stained sections of knee joints from 12-month male control **(A)** and male Mig-6 over **(B)** mice were evaluated for cartilage damage following OARSI histopathological scale on the two quadrants of the knee: MFC = medial femoral condyle, MTP = medial tibial plateau. OARSI based cartilage degeneration scores are significantly higher in the MFC and MTP of Mig-6 overexpressing mice, corresponding to the increased damage observed histologically **(C)**. Data analyzed by two-way ANOVA with Bonferroni’s multiple comparisons test. Individual data points presented with mean ± SEM. All scale bars =100 μm. N = 9 mice/group.

**Figure 6).**
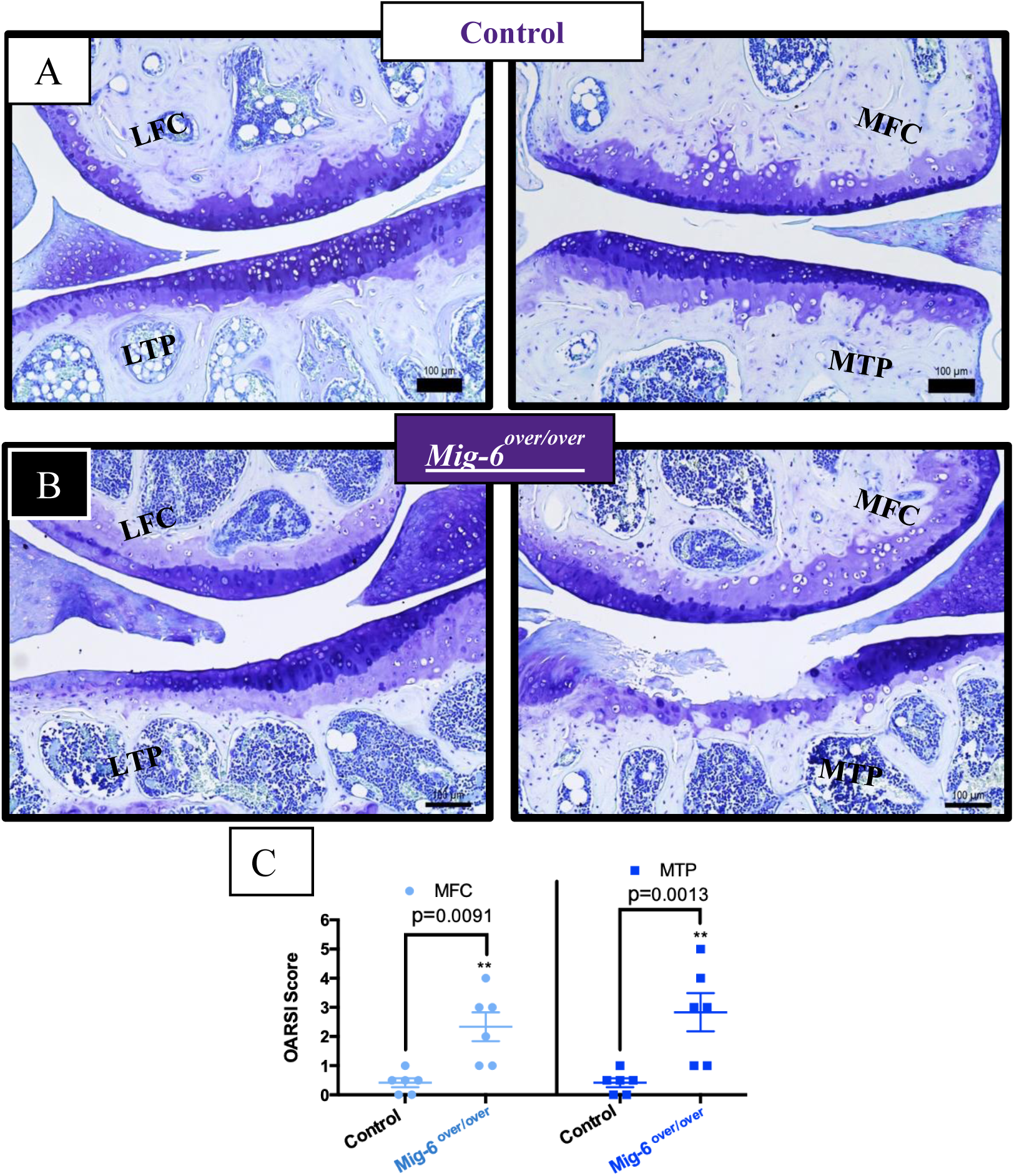
18 months old *Mig-6*^*over/over*^ mice leads to advanced OA-like cartilage. Representative images of Toluidine Blue stained sections of knee joints from 18-month male control **(A)** and male Mig-6 over **(B)** mice were evaluated for cartilage damage following OARSI histopathological scale on the two quadrants of the knee: MFC = medial femoral condyle, MTP = medial tibial plateau. OARSI based cartilage degeneration scores are higher both in the MFC and MTP of Mig-6 overexpressing mice, corresponding to the increased damage observed histologically. **(C)** Data analyzed by two-way ANOVA with Bonferroni’s multiple comparisons test. Individual data points presented with mean ± SEM. All scale bars =100 μm. N = 6 mice/group.

### Overexpression of *Mig-6* decreases EGFR phosphorylation and Sox9 expression

Since Mig-6 negatively regulates EGFR signaling (32,33,41), immunohistochemistry was performed for phospho-EGFR (Tyr-1173) (pEGFR), with no primary antibody controls. Frontal knee sections from 11 weeks-old male *Mig-6* ^*over/over*^ mice showed decreased pEGFR staining in the medial compartment in the knee joint (Fig.7), as expected upon Mig-6 overexpression.

During chondrogenesis, the transcription factor SOX9 is required for cartilage formation and normal expression of collagen and aggrecan (51). Sagittal and frontal sections of paraffin embedded knees from post-natal day 0 (P0), 6 weeks-old, 11 weeks-old, 12 months and 18 months male mice were used for SOX9 immunostaining. At P0, nuclear SOX9 expression was observed in the resting and proliferative zone of the growth plate in both genotypes (Fig. 8A,B). Cell density was not different between genotypes (Fig. 8C). In control mice, 78 % of chondrocytes were positive for SOX 9 immunostaining, while the proportion of positive cells was only 53 % in *Mig-6* ^*over/over*^ mice (Fig 8D). In 6 and 11-weeks-old mice, SOX9 was present in the articular cartilage in all four quadrants (Fig. 9A,B). At 6 weeks-old the total cell number in control male and *Mig-6*^*over/over*^ mice is similar (Fig. 9C), but the percentage of SOX9 positive cells was decreased in mutant mice (Fig 9D). A similar phenotype was present at 11 weeks (Supplementary Fig 7). At 12 months of age, SOX9 is present more in the lateral side (LTP and LFC) than the medial side (MTP and MFC) in both strains, with a few positive cells present in the medial side of the control strain. On the other hand, *Mig-6* ^*over/over*^ mice showed fewer SOX9-positive cells on the medial side due the articular cartilage damage (Fig 10). Similar results were found at 18 month of age in *Mig-6* ^*over/over*^ with decreased SOX9 immunostaining in their medial side compared to the control (data not shown). For all ages, negative controls did not show staining in chondrocytes.

**Figure 7).**
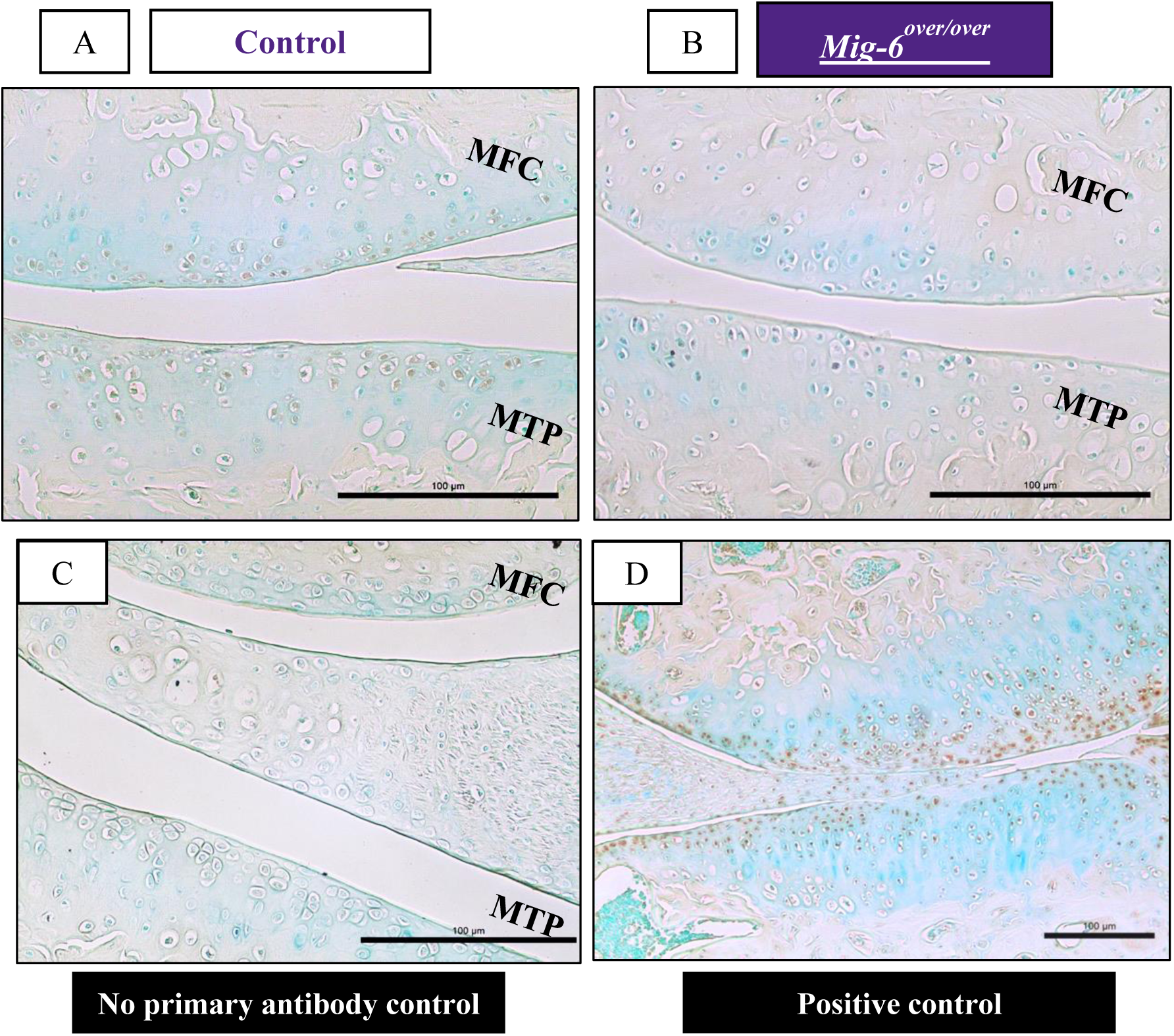
Phospho-EGFR staining is decreased in the articular cartilage of cartilage specific Mig-6 overexpressing mice at 11 weeks of age. Immunostaining of phosphorylated epidermal growth factor receptor (pEGFR; Tyr-1173) in the knee joints of 11 week old *Mig-6* ^*over/over*^*Col2a1-Cre*^*+/-*^ **(B)** is decreased in response to increased Mig-6 levels. Frontal sections of mice articular cartilage, as negative control, exhibited no staining **(C)**. Also, cartilage-specific deletion of *Mig-6*, serving as positive control **(D)**. N=5 mice/genotyping. MFC = medial femoral condyle and MTP = medial tibial plateau. Scale bar = 100µm.

**Figure 8).**
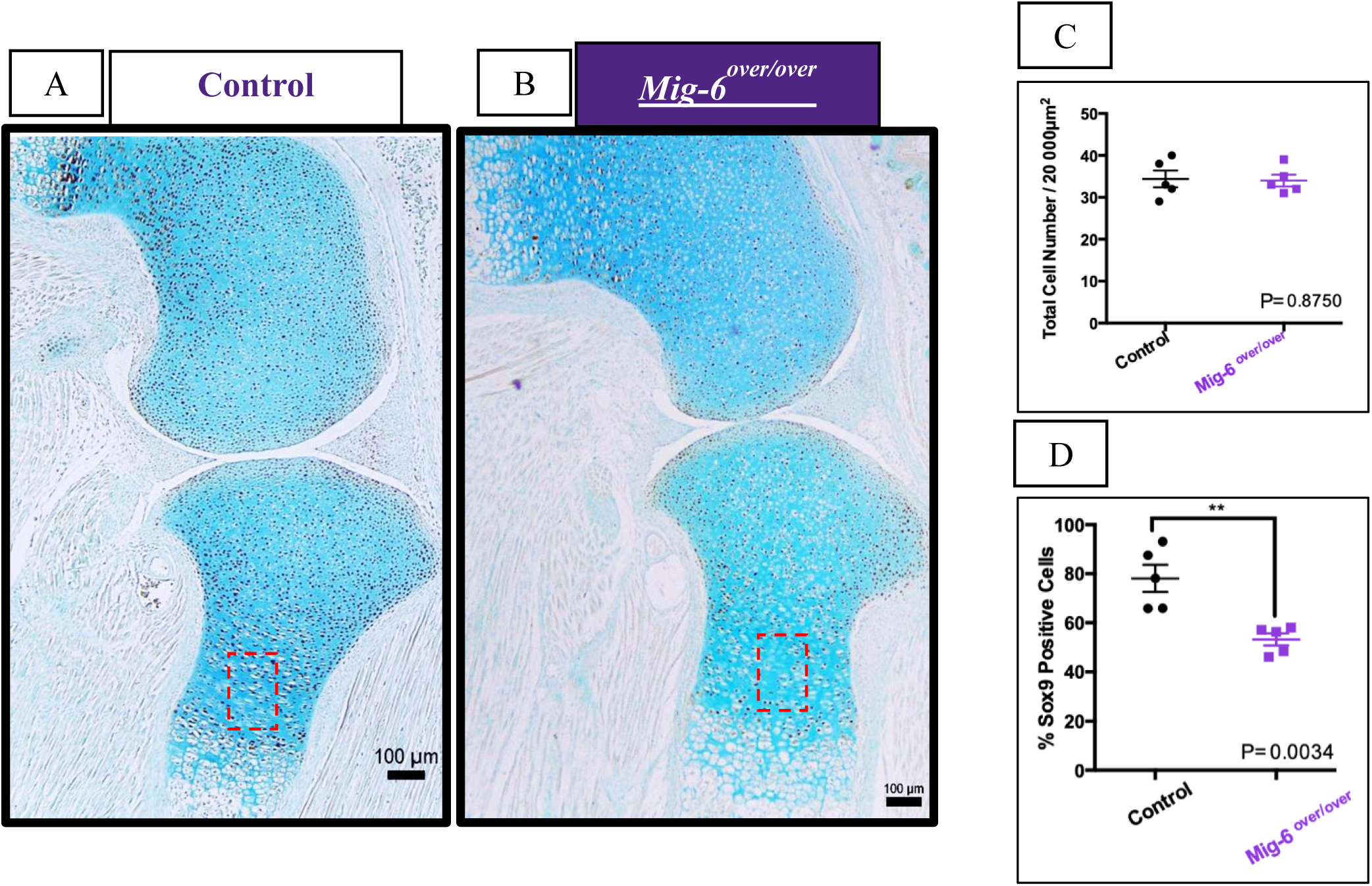
SOX9 immunostaining shows a decrease in Mig-6 overexpressors mice at post natal day 0 (p0). Ratio between the total cell number from control and Mig-6over **(B).** Ratio between the percentage of Sox9 positive cells from control and Mig-6over at p0 mice **(C).** Data analyzed by two tailed student t-tests from 5 mice per group. Individual data points presented with mean ± SEM (P<0.05). Scale bar = 100µm.

**Figure 9).**
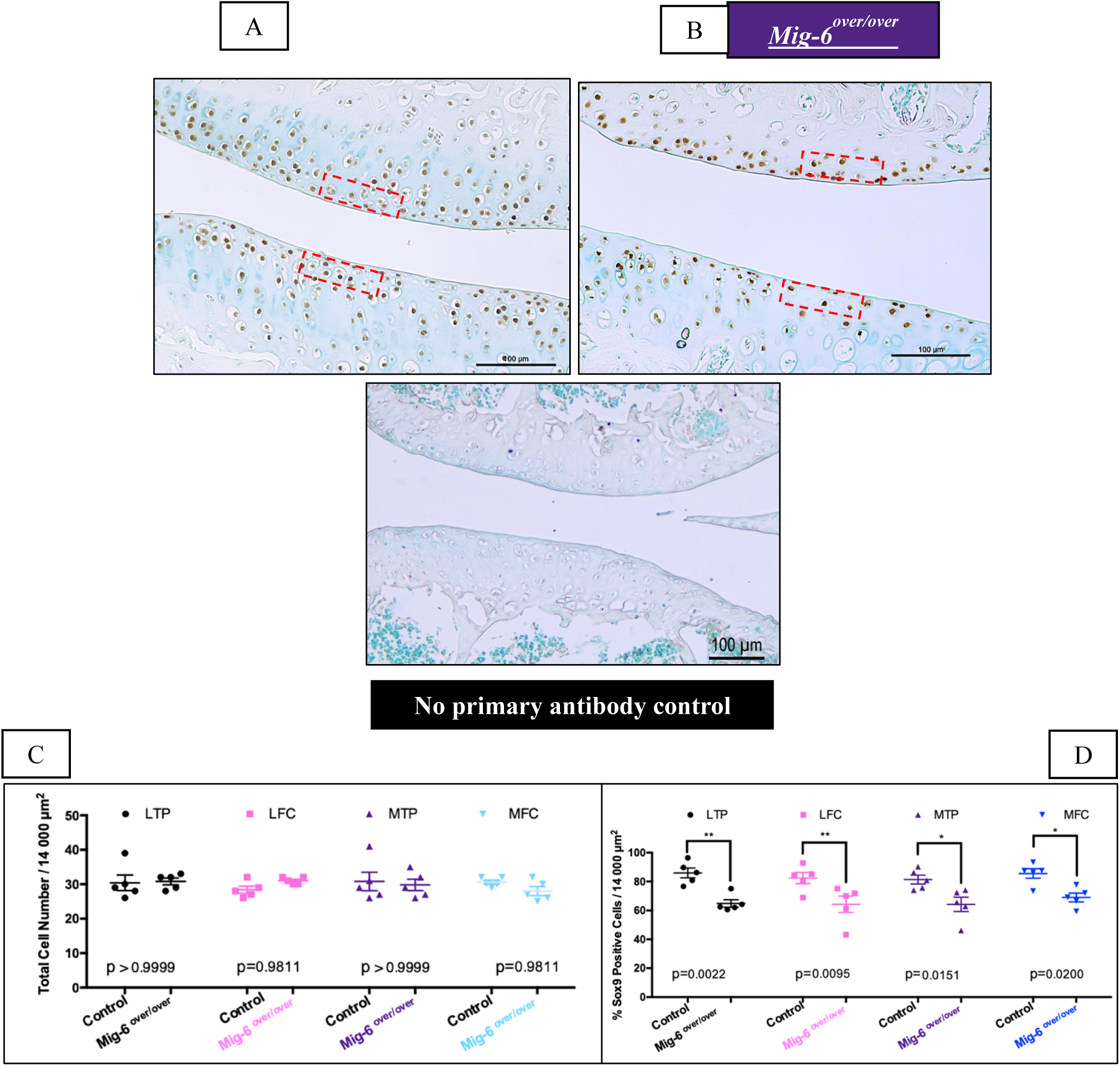
SOX9 immunostaining shows a decrease in Mig-6 overexpressors mice at 6 weeks-old male mice control and Mig-6over. No primary antibody for SOX9 display no staining with methyl green countersain in mice. Ratio between the total cell number from control and Mig-6over at 6 weeks-old male mice **(C).** Ratio between the percentage of Sox9 positive cells from control and Mig-6over at 6 weeks-old male mice **(D).** Data analyzed by two-way ANOVA (95% CI) with Bonferroni post-hoc test. Individual data points presented with mean ± SEM; N= 5 mice/genotyping. LFC = lateral femoral condyle, LTP = lateral tibial plateau, MFC = medial femoral condyle and MTP = medial tibial plateau. Scale bar = 100µm.

**Figure 10).**
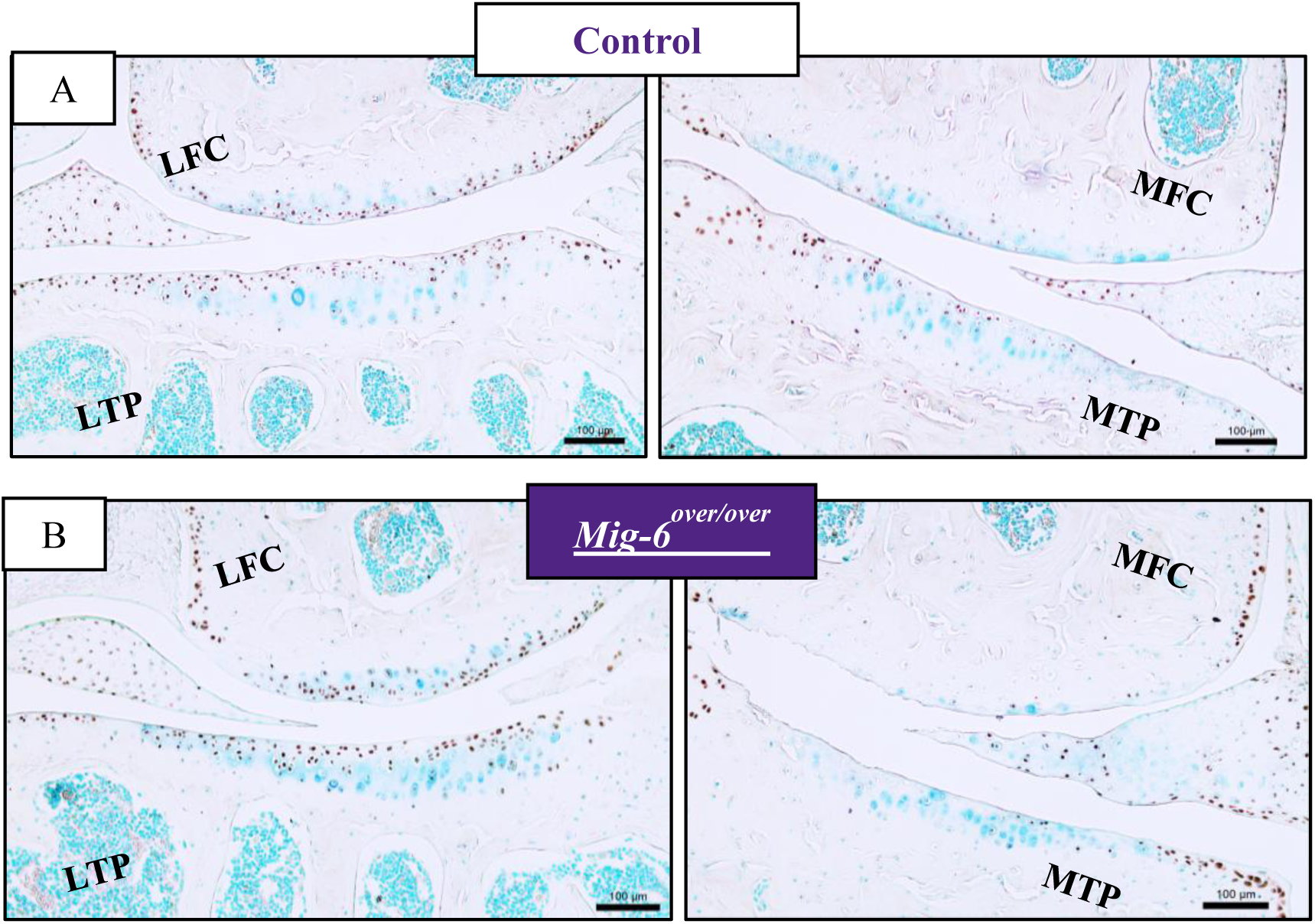
12-month-old cartilage specific Mig-6 overexpressing mice show decreased SOX9 immunostaining. Representative SOX9 immunostained in male mice (n=5) in MFC and MTP show decreased staining intensity in Mig-6 over mice **(B)** when compared to control **(A).** No primary control for articular cartilage **(C).** LFC = lateral femoral condyle, LTP = lateral tibial plateau, MFC = medial femoral condyle and MTP = medial tibial plateau. Scale bar = 100µm.

### Overexpression of Mig-6 decreases expression of lubricin

Lubricin (aka PRG4/superficial zone protein) is a proteoglycan that plays an important role as lubricant in the joint (52). EGFR signaling is crucial for the cartilage lubrication function and regulates the induction of *Prg4* expression which is necessary for smooth movement (33,53). Immunohistochemistry for Lubricin in 11 week-old and 12 months-old animals demonstrated less staining in the superficial zone of the medial side of *Mig-6* ^*over/over*^ mice than in the control group (Fig.11 and Fig. 12).

**Figure 11).**
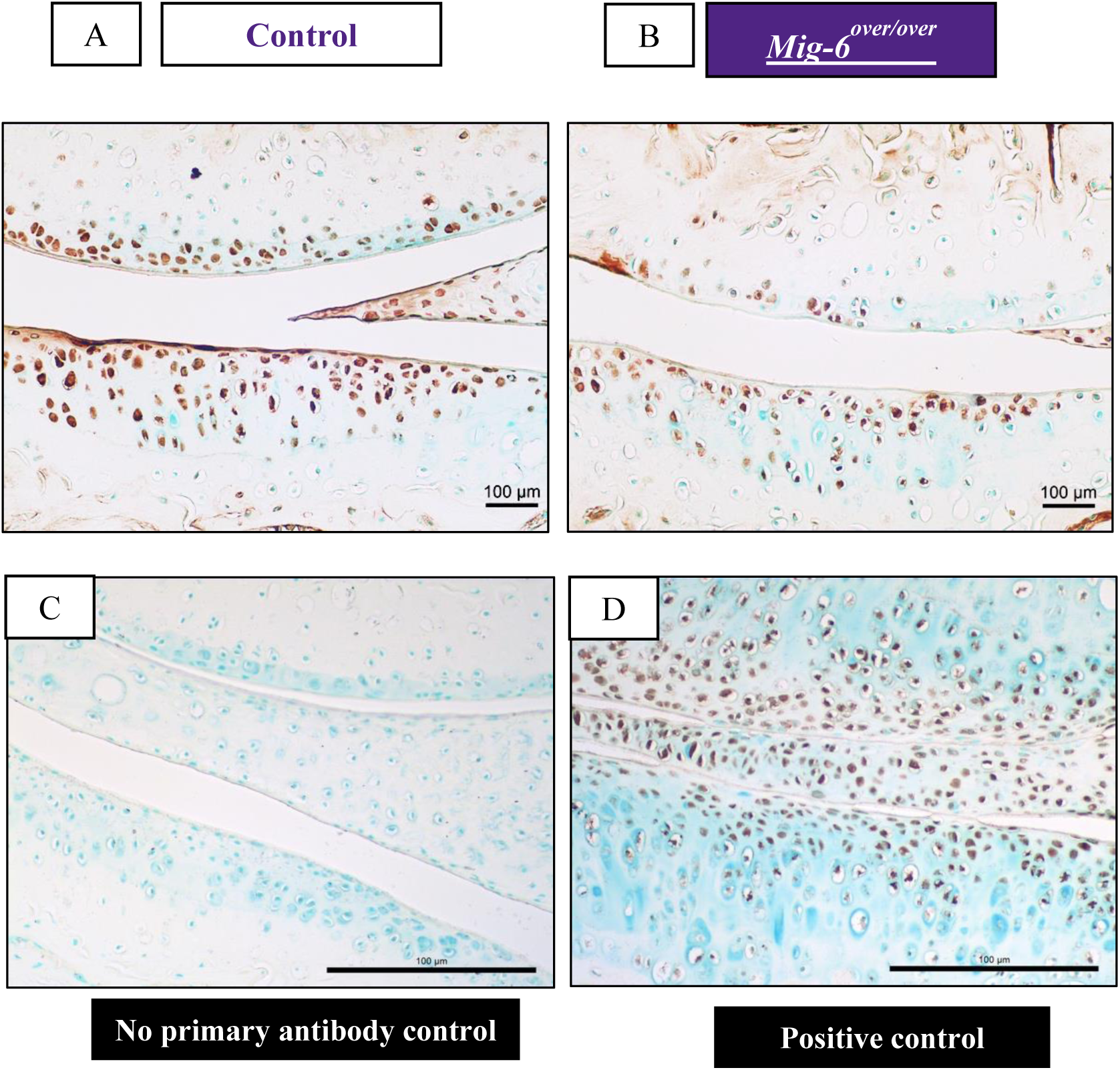
Lubricin immunostaining is slightly decreased in the articular cartilage of cartilage specific Mig-6 overexpressing mice at 11 weeks of age. Immunostaining of sections of the knee joint indicate the presence of Lubricin (*PRG4*) in the superficial zone chondrocytes. IHC reveals no staining for the negative control **(C)** and *Mig-6* KO, serving as positive control **(D)**. N=4-5 mice/genotyping. MFC = medial femoral condyle and MTP = medial tibial plateau. Scale bar = 100µm.

**Figure 12).**
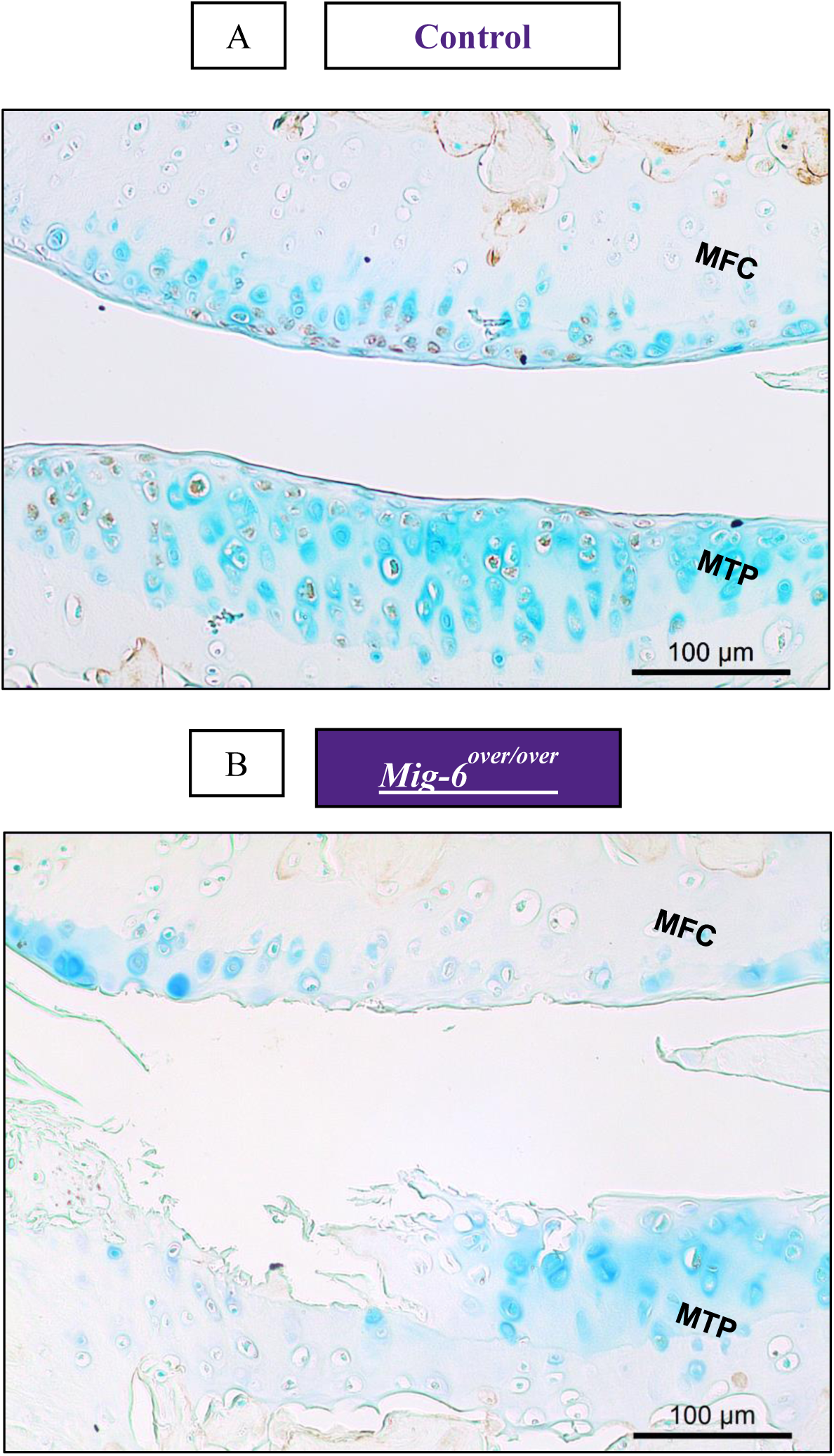
Lubricin immunostaining is decreased in the articular cartilage of cartilage specific Mig-6 overexpressing mice at 12 months of age. Immunostaining of sections of the knee joint indicate the presence of Lubricin (*PRG4*) in the superficial zone chondrocytes. N=4-5 mice/genotyping. MFC = medial femoral condyle and MTP = medial tibial plateau. Scale bar = 100µm.

### MMP13 immunostaining is similar in Mig-6-overexpressing and control mice

Matrix metalloproteinase (MMP) 13 is highly expressed in OA (54,55). Frontal sections of knees from 12- and 18-month-old control and *Mig-6* ^*over/over*^ male mice were used for MMP13 immunohistochemistry. At 12 months, pericellular staining was observed in the lateral articular cartilage of male mice from both genotypes, along with the expected subchondral bone staining (Fig. 13). Less staining was observed on the medial side of control mice while advanced cartilage degeneration in mutant mice precluded staining. Negative controls did not show staining in cartilage or subchondral bone. Articular cartilage from 18 months-old mice showed similar staining patterns and intensity of MMP13 immunostaining in the lateral side of both genotypes, however in the medial side of *Mig-6* ^*over/over*^ mice, MMP13 staining is seen on the cartilage surface (lesion sites) and also observed in the subchondral bone (data not shown).

**Figure 13).**
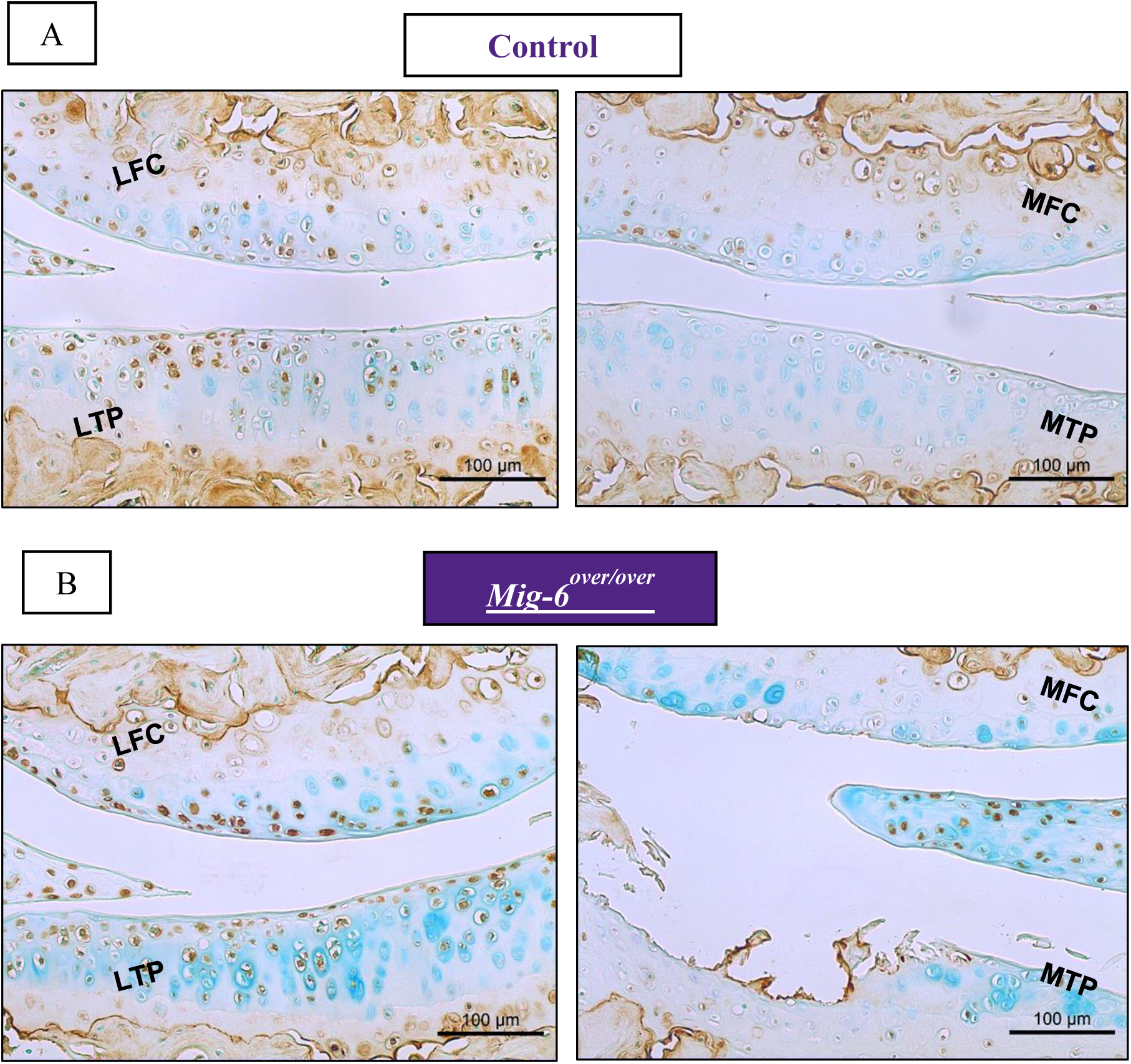
12 month-old cartilage specific Mig-6 overexpressing mice show similar pattern of MMP13 as the control mice. Representative immunohistochemistry of matrix metalloproteinase 13 (MMP13) for Mig-6 overexpression mice at 12 months control **(A)** and cartilage specific Mig-6 overexpression **(B).** No primary control for articular cartilage **(C).** N=5 mice/genotyping. LFC = lateral femoral condyle, LTP = lateral tibial plateau, MFC = medial femoral condyle and MTP = medial tibial plateau. Scale bar = 100µm.

## Discussion

The maintenance of articular cartilage homeostasis relies on a dynamic equilibrium involving growth factors (56), genetics (57), mechanical forces (58), obesity and injury, that all play a role in the onset of osteoarthritis (59). Better understanding of the underlying molecular mechanism is required to design therapies for preventing progression of OA. Recent studies from our laboratory and others have identified the epidermal growth factor receptor (EGFR) and Mig-6 as possible mediators of articular cartilage homeostasis (35,41,53,60). *Mig-6* is a cytosolic protein and negative feedback regulator of EGFR signalling (61); thus, *Mig-6* can be a potential tumor suppressor (43,62–65). In addition, whole body knockout of the *Mig-6* gene in mice results in degenerative joint disease (38). We also have shown previously that constitutive cartilage-specific deletion of *Mig-6* (Mig-6 KO) results in increased articular cartilage thickness and cell density in the joints of 12 week-old mice (41). Cartilage-specific Mig-6 KO mice show the same anabolic effect in joint cartilage at 21 months of age (unpublished).

Previous research demonstrates that Mig-6 overexpression acts as a negative feedback regulator of EGFR-ERK signalling (43), however these studies did not yet analyze joint tissues. Since our studies suggest dosage- and/or context-specific roles of EGFR signaling in joint homeostasis and OA (35), we now examined whether overexpression of Mig-6 alters these processes. Here, we report that cartilage-specific constitutive overexpression of *Mig-6* did not cause cartilage degeneration in young mice, but early onset OA in middle aged mice. While we observed some effects of Mig-6 overexpression on bone length and weight, these effects were subtle and not accompanied by major morphological or histological changes in growth plate cartilage, overall skeletal morphology, or body composition. A previous study showed that deletion of EGFR in bone tissue (*Col1-Cre Egfr*^*Wa5/f*^) resulted in shorter femurs compared to wild-type mice (66), consistent with our findings. The EGFR network is essential during long bone development, since previous studies have shown that EGFR- or TGFα-deficient mice exhibit a widened zone of hypertrophic chondrocytes (24,67). Moreover, Qin and colleagues have shown that administration of the EGFR inhibitor, gefitinib, into 1-monht-old rats results in an enlarged hypertrophic zone due down-regulation of MMP-9,-13 and -14 (31). Together these data suggest a critical role of EGFR during endochondral ossification and elucidate downstream mechanism of EGFR (68). Further research is required to provide more evidence of EGFR/*Mig-6*^*over/over*^ signalling during bone formation, but many of these effects are relatively subtle and transient, and likely unrelated to much more severe phenotypes observed later.

Histologically, our findings showed that mice with cartilage-specific *Mig-6* overexpressing showed healthy articular cartilage with no significant difference in articular cartilage thickness from control group at the ages of 6 weeks and 11 weeks. However, *Mig-6* ^*over/over*^ mice developed severe degeneration of articular cartilage with aging. More prevalent, the knee joints of *Mig-6* ^*over/over*^ male mice showed significantly advanced cartilage degeneration. The same pattern but with more severe damage, was seen in 18 month-old mice. As previously described, sex hormones play a role in OA disease where male mice develop more severe OA (69).

SOX9 is crucial in chondrogenesis during endochondral bone formation, articular cartilage development and cartilage homeostasis (51). Previous *in vivo* models using cartilage (Col2)-Cre or limb (Prx1)-Cre specific ablation of *Mig-6* showed increased expression of SOX9 in the articular cartilage. Also, TGFα supresses expression of anabolic genes such as Sox9, type II collagen and aggrecan in primary chondrocytes (71). Interestingly, in the medial and lateral compartment of the knee joints of 6 and 11 week-old male *Mig-6* ^*over/over*^ mice, the percentage of SOX9-positive chondrocytes was decreased compared to controls, despite the absence of histological defects in articular cartilage. These data suggest that reduced number of Sox9-expressing cells precede the degeneration of articular chondrocytes in our mutant mice. The number of SOX9-expressing cells was also reduced in *Mig-6* ^*over/over*^ mice at later ages. These data suggest that reduced numbers of Sox9-expressing cells could be one cause of the advanced OA in our mutant mice. In addition, we observed decreased expression of lubricin/PRG4 in these joints, which might also contribute to the observed joint pathologies. PRG4 has been shown to be regulated by EGFR signaling before (42,53), in support of our findings.

While the EGFR is the best characterized substrate of Mig-6, other substrates have been described. Mig-6 binds to different proteins such as the cell division control protein 42 homolog (Cdc42) (72), c-Abl (73), and the hepatocyte growth factor receptor c-Met (74). While we cannot exclude that deregulation of these other substrates contributes to the observed phenotypes, the similarities of defects in our mice with those seen upon cartilage-specific deletion of EGFR suggest that decreased EGFR signaling is the main cause for the advanced OA observed in our mutant mice. Nevertheless, it will be important to determine whether signaling through cMet and other pathways is altered as well.

In conclusion, we show for the first time that cartilage-specific Mig-6 overexpression in mice results in reduced EGFR activity in chondrocytes, reduced SOX9 and PRG4 expression, and accelerated development of OA. These data highlight the important and context-specific role of the EGFR-Mig-6 signaling pathway in joint homeostasis and point towards potential targeting of this pathway for OA therapy.

## Acknowledgements

We would like to thank Julia Bowering for kindly performing articular cartilage sectioning and Dr. Michael Pest for his assistance in the blinded scoring of joints. M.B. was supported by a fellowship from CNPq/Brazil. Work in the lab of F.B. is supported by a grant from the Canadian Institutes of Health Research (Grant #332438). F.B. holds the Canada Research Chair in Musculoskeletal Research.

