## Supplementary Figures for "Overexpression of mig-6 in cartilage induces an osteoarthritis-like phenotype in mice"

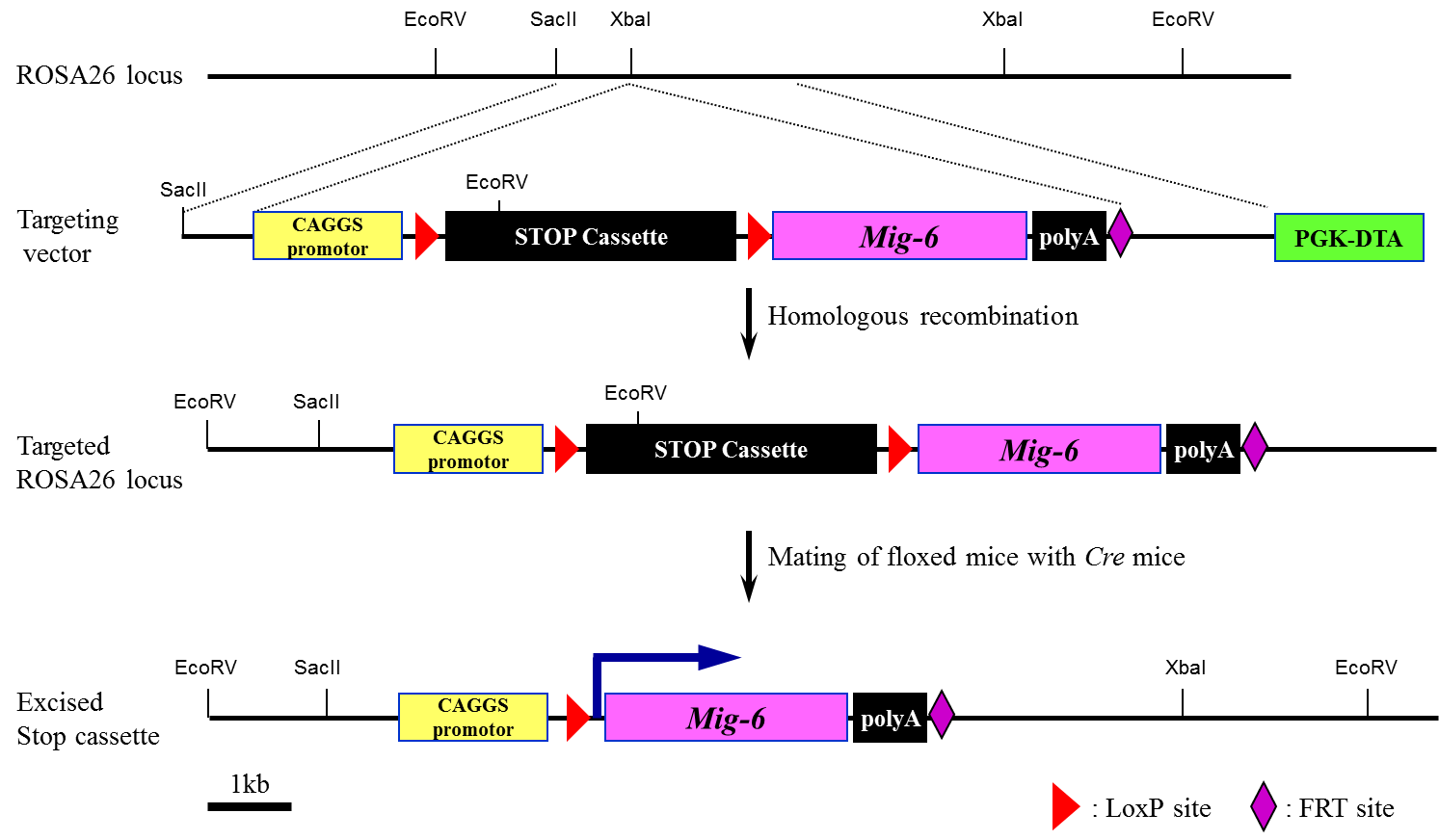

**A**

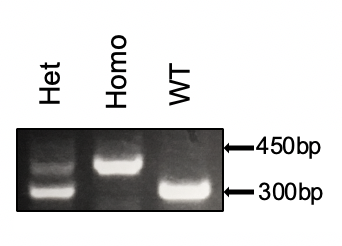

**C**

**B**

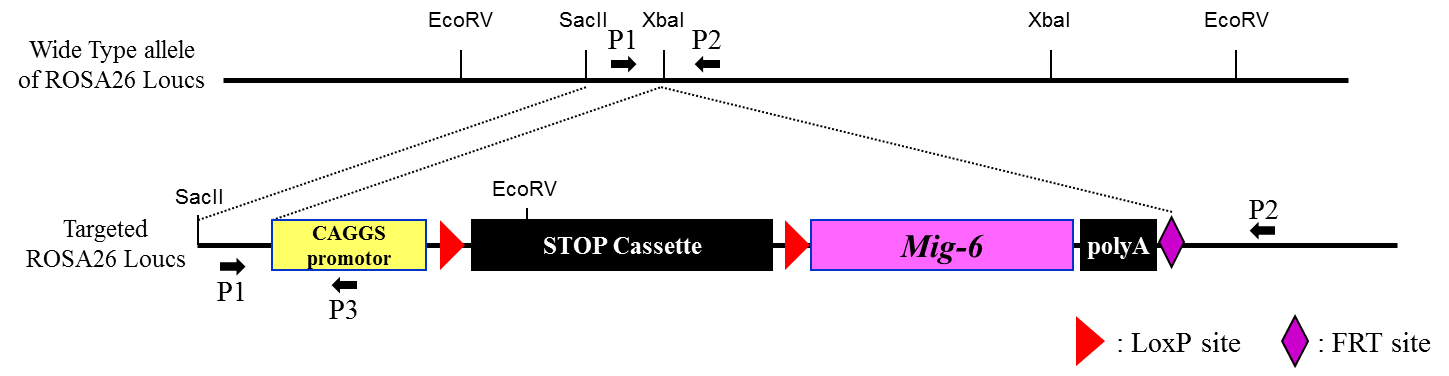

**D**

| PCR Primer | 5’ to 3’ |
| --- | --- |
| P1 | GGG GGA GGG GAG TGT TGC |
| P2 | CCA TTT TCC TTA TTT GCC CCT ATT |
| P3 | GGG CCA TTT ACC GTC ATT |

**Supplementary Figure 1) Construction of the targeting vectors and generation of *Mig-6^over/over^*** **mice.** Adapted from Kim, T. H. *et al.* Mig-6 suppresses endometrial cancer associated with pten deficiency and ERK activation. *Cancer Res.* **74,** 7371–7382 (2014). **(A)** The overexpression of *Mig-6* is accomplished by placing the transcription of *Mig-6* under the control of a ubiquitously expressed promoter, the chicken b actin-cytomegalovirus hybrid (CAGGS) promoter. The construction also contained the “Stop Cassette” flanked by LoxP sites (LSL). **(B)** PCR strategy. P1 and P2 can amplify a 300 bp fragment from the wild-type allele, whereas P1 and P3 can amplify a 450 bp fragment from the targeted ROSA26 locus allele. **(C)** A representative agarose gel image of PCR genotyping, heterozygous (Het), wild-type (wt) and homozygous (Homo). **(D)** PCR primer sequence for wild type and *Mig-6^LSL^* allele.

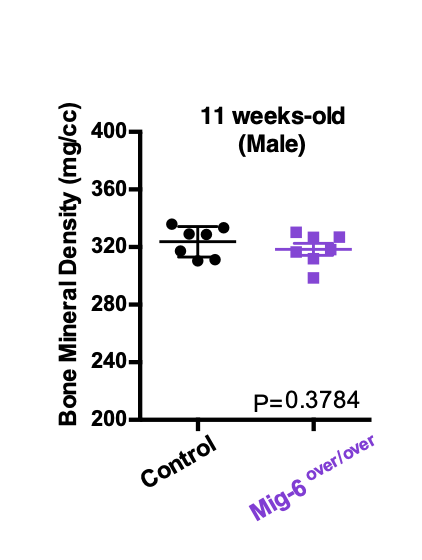

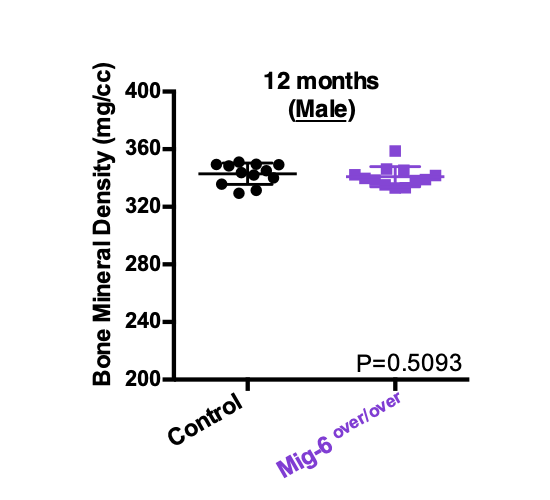

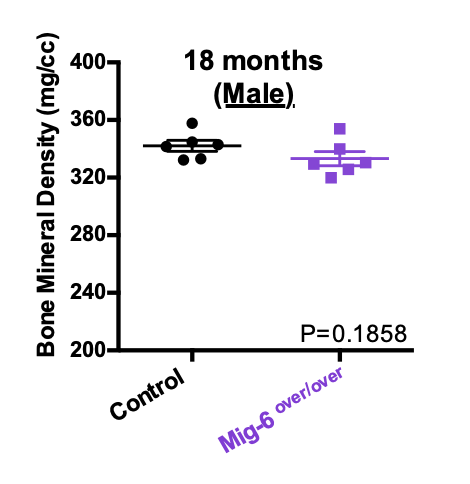

**C**

**B**

**A**

**Supplementary Figure 2) Bone mineral densities were measured from µCT scan volumes from control and *Mig-6^over/over^*** **male mice. (A)** Mean bone mineral densities were 323.7 mg/cc (control) and 318.5 mg/cc (*Mig-6^over/over^*) at 11 weeks-old. Moreover, **(B)** At 12 months of age male *Mig-6^over/over^* mice had mean bone mineral density (342.9 mg/cc) and controls male mice (341.0 mg/cc). **(C)** At 18 months of age, there were no significance difference between the mean bone mineral density from control male mice (342.1 mg/cc) and male *Mig-6^over/over^* (333.2 mg/cc). There were not significantly different among 11 weeks-old, 12 and 18 months for bone mineral density for male control and Mig-6 overexpression mice. Individual data points presented with mean ± SEM (P<0.05). Data analyzed by two tailed student t-tests from 6-12 mice per group (age/gender).

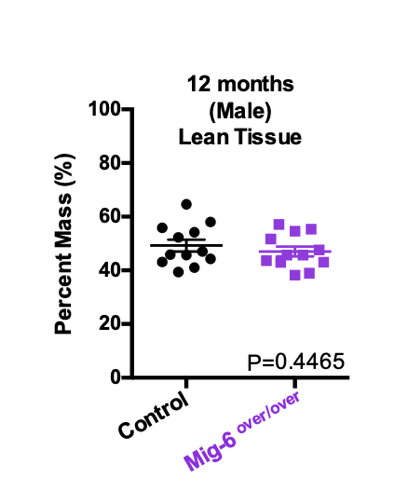

**B1**

**A1**

**A2**

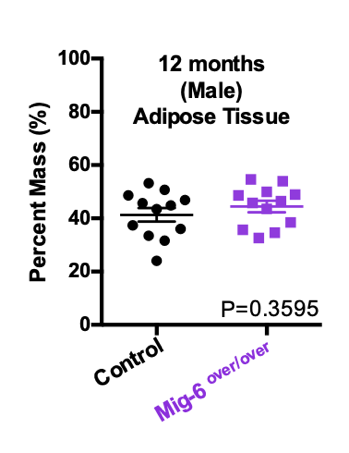

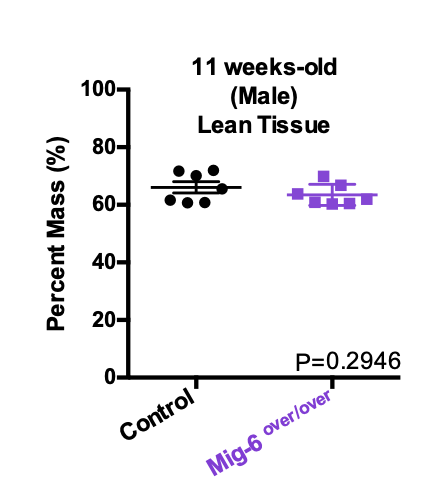

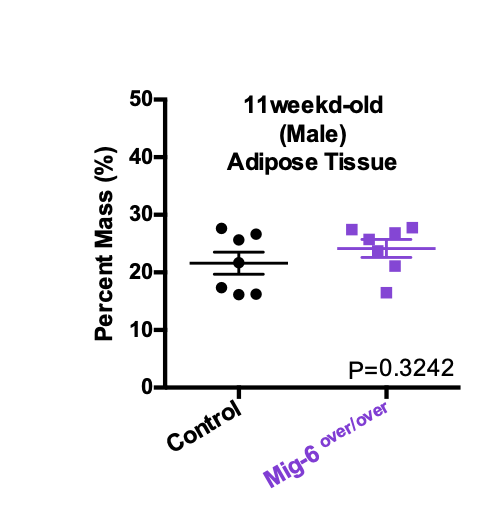

**C1**

**C2**

**B2**

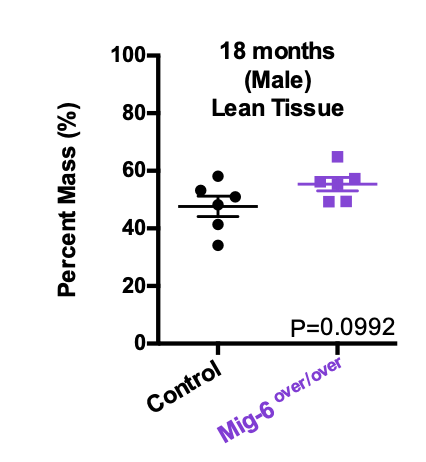

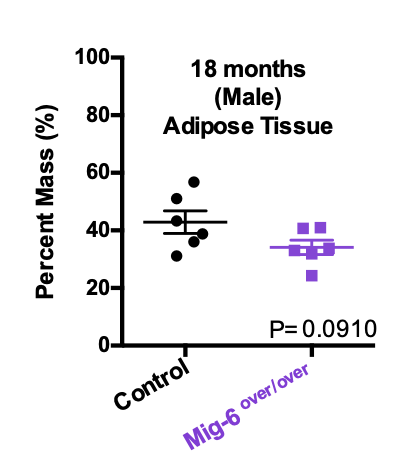

**Supplementary Figure 3) Body composition and body mass from growth and aging male control and *Mig-6^over/over^***^.^ Body composition was calculated from control and *Mig-6^over/over^* male mice. At 11 weeks-old **(A1/A2)**, 12 months **(B1/B2)** and 18 months **(C1/C2)** neither the average lean mass percent or mean body fat were not statistically significant between genotypes. Individual data points presented with mean ± SEM (P<0.05). Data analyzed by two tailed student t-tests from 6-12 mice per group (age/genotyping).

**B**

**Mig6 ^over/over^**

**A**

**Control**

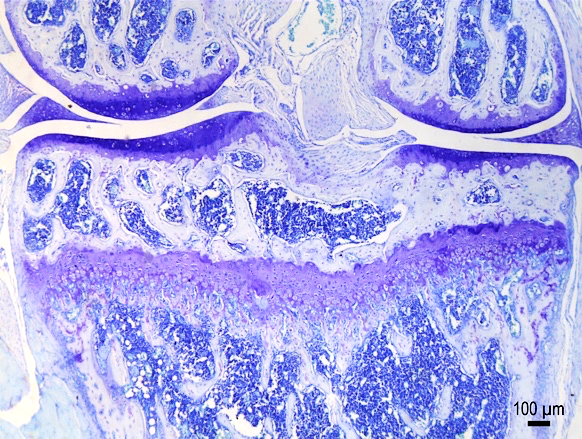

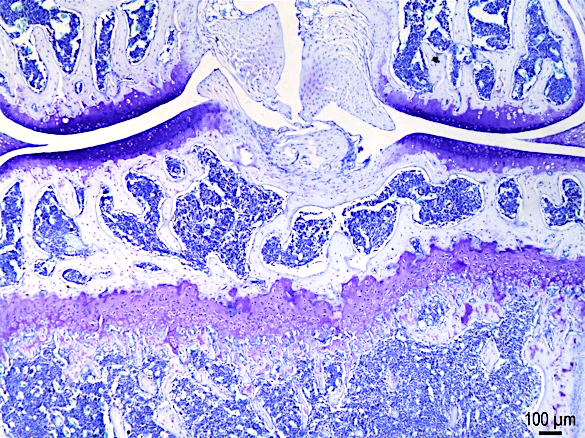

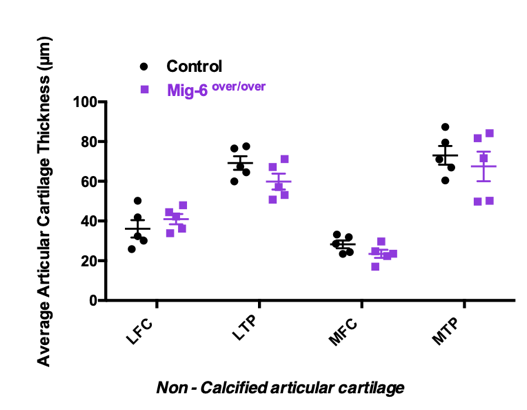

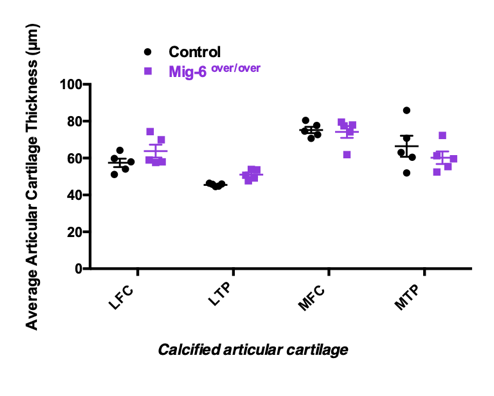

**D**

**C**

**Supplementary Figure 4)** **Articular cartilage from 11 weeks-old *Mig-6^over/over^*** **female mice appeared healthy during skeletal maturity.** Representative (n=5/group, toluidine blue) stained frontal sections of knee joints from 11-week-old control **(A)** and Mig-6over **(B).** Mig-6 overexpressors mice show similar articular cartilage thickness when compared to controls at 11 weeks-old female mice. The average thickness of the calcified articular cartilage **(C)** and non-calcified articular cartilage **(D)** in the lateral femoral condyle (LFC), lateral tibial plateau (LTP), medial femoral condyle (MFC), medial tibial plateau (MTP) was measured. Individual data points presented with mean ± SEM. Data analyzed by two-way ANOVA (95% CI) with Bonferroni post-hoc test. Scale bar = 100µm.

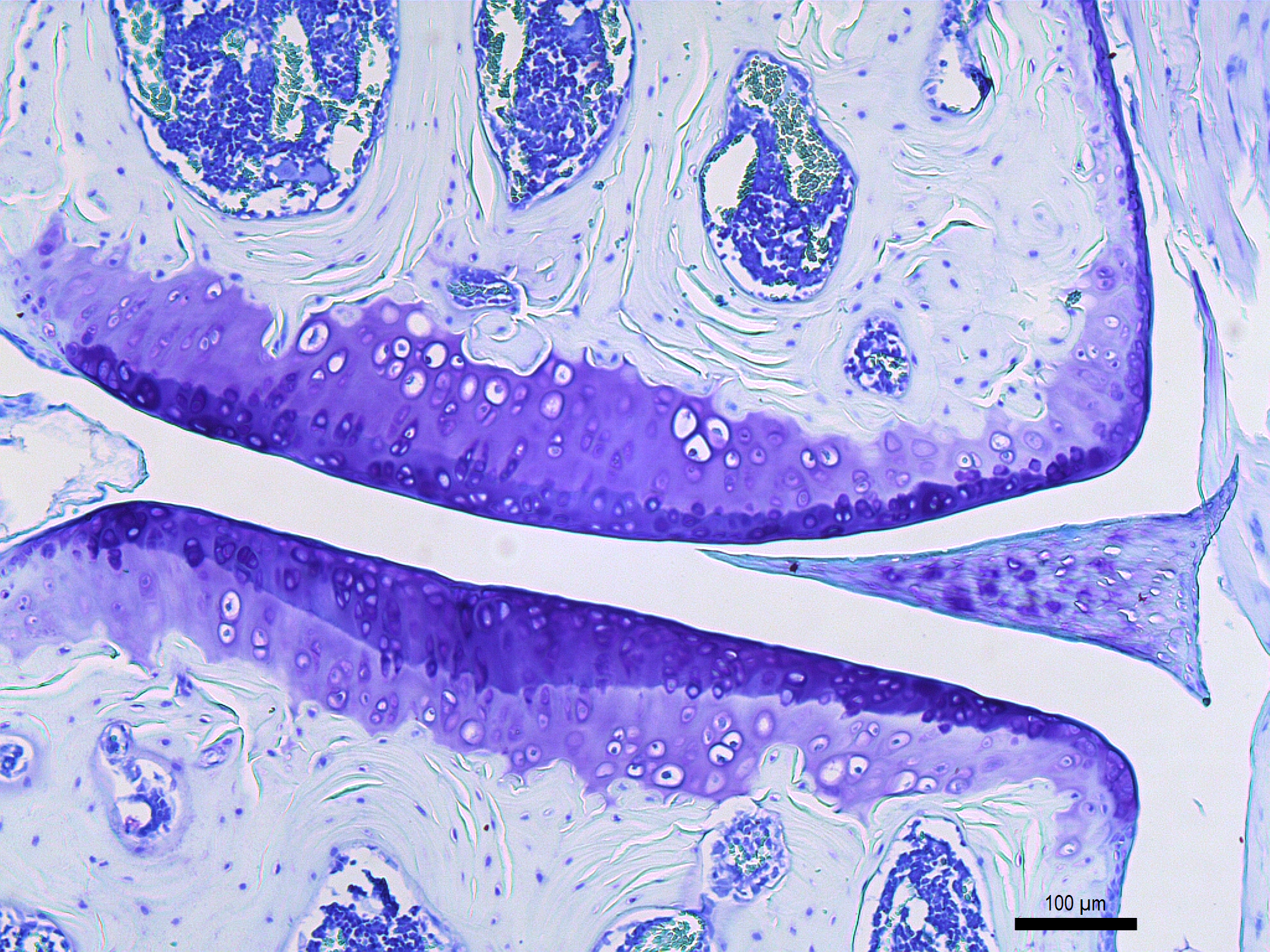

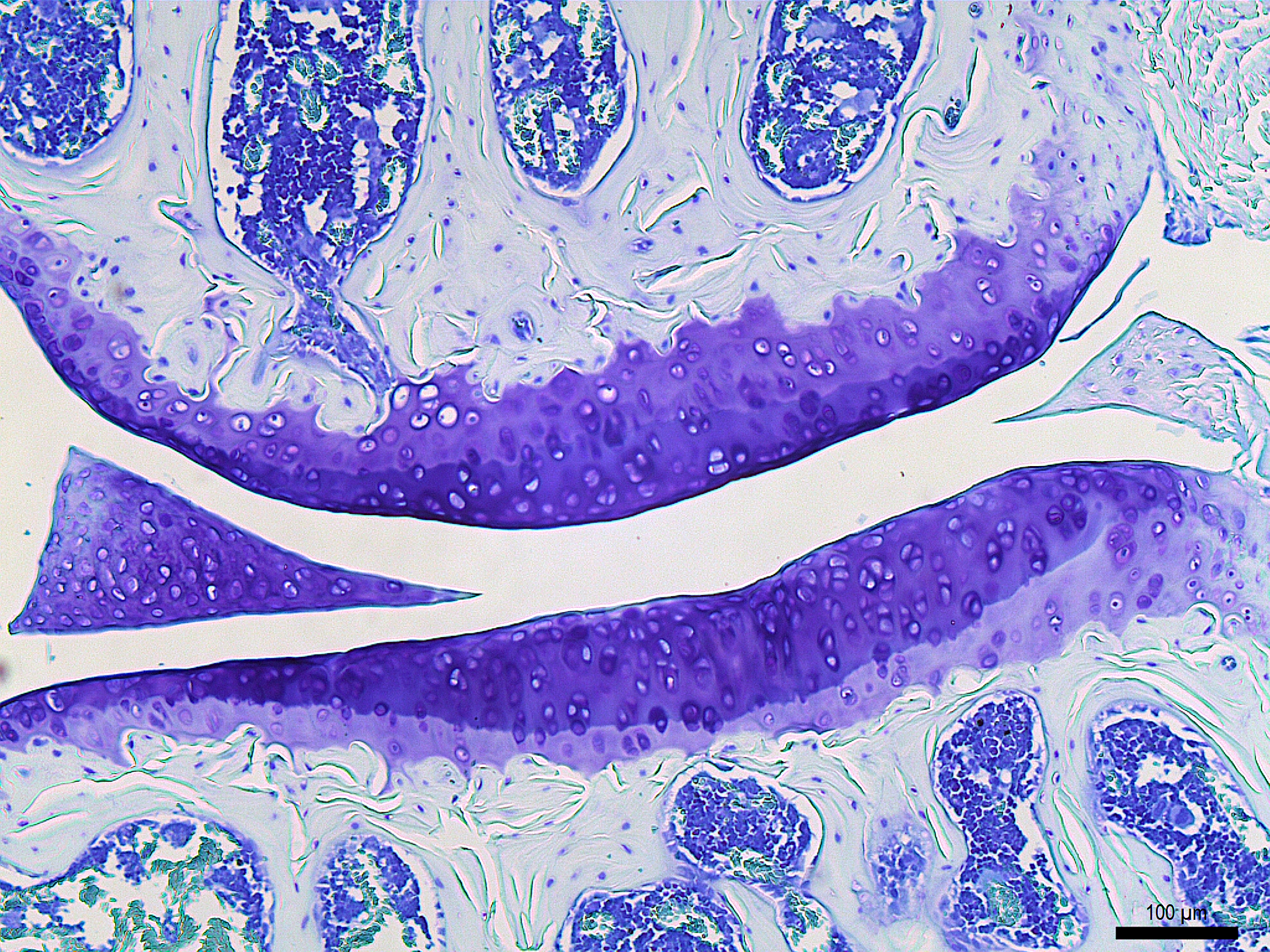

C

**12 months old**

**MFC**

**Control**

A

**LFC**

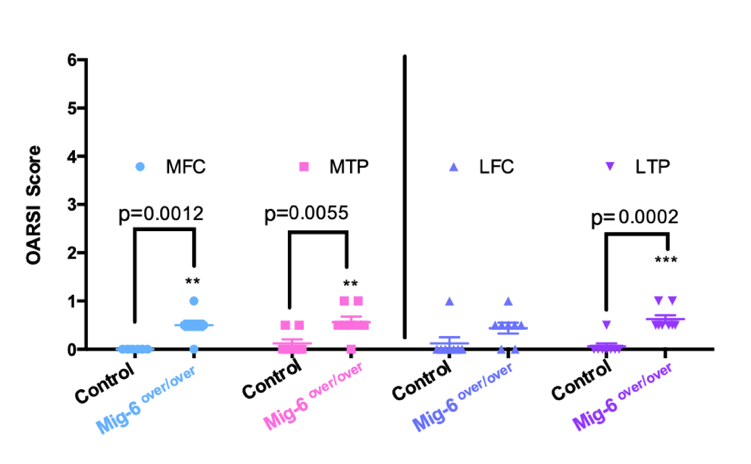

**MTP**

**LTP**

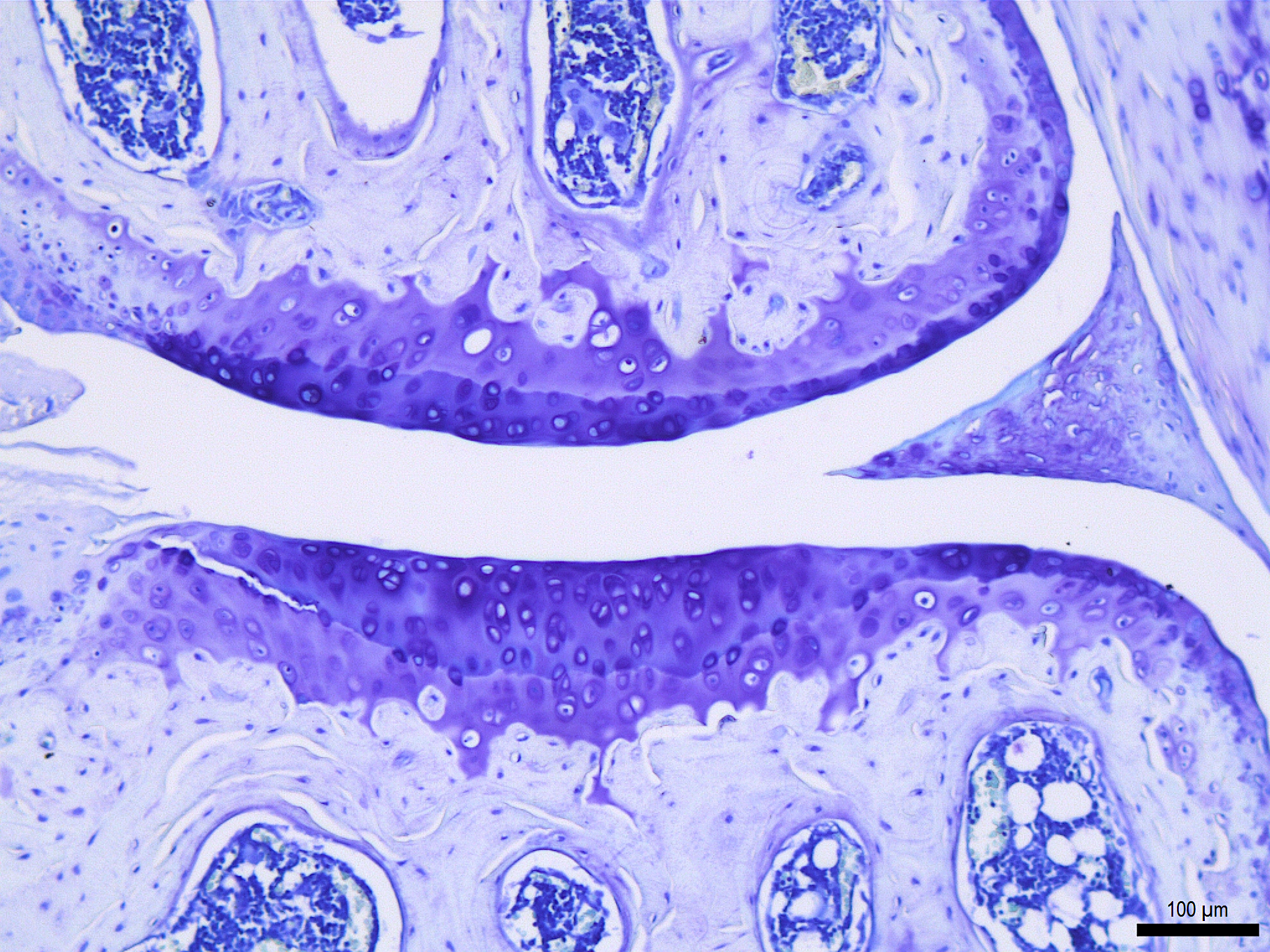

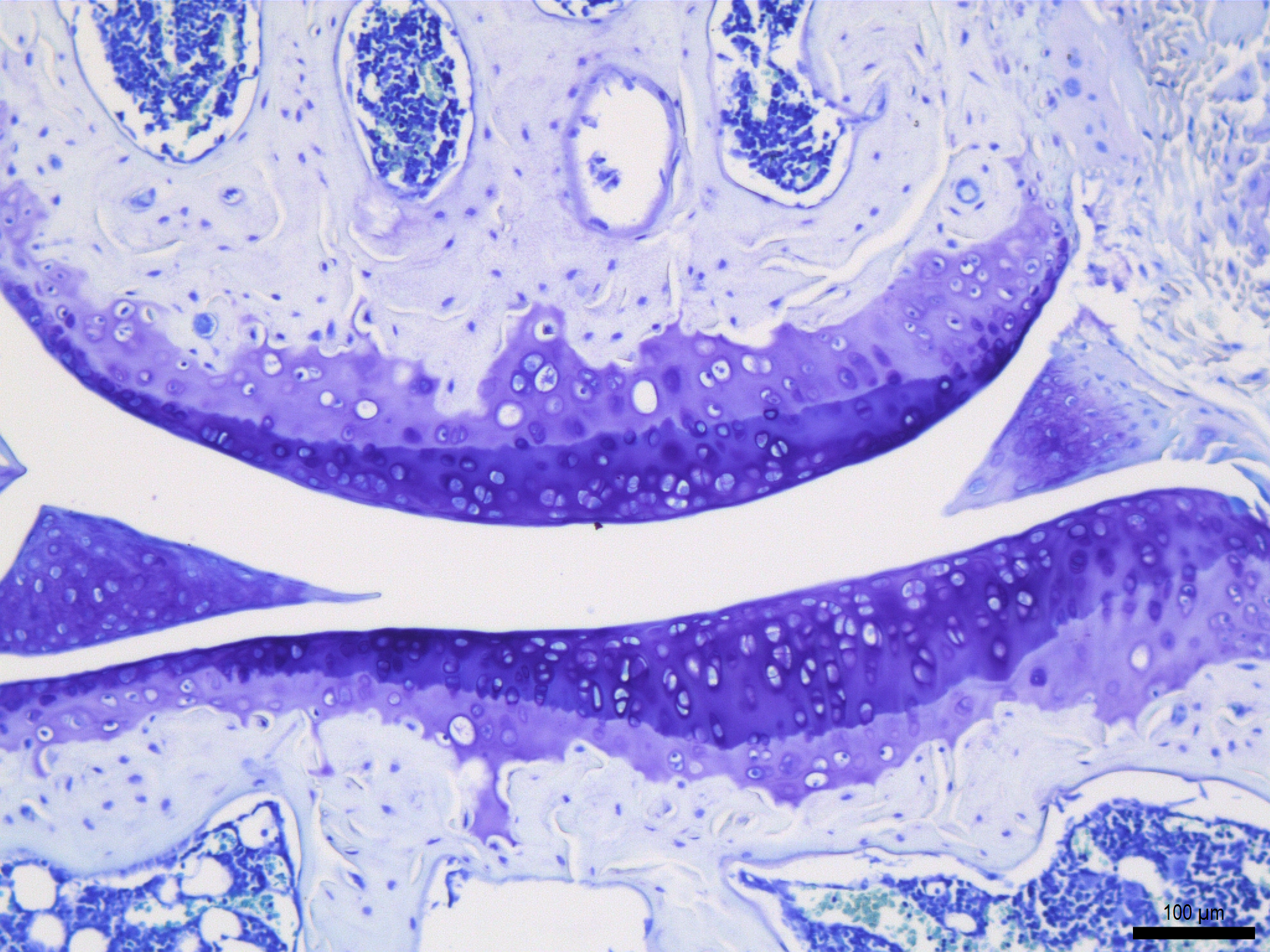

B

***Mig-6^over/over^***

**MFC**

**LFC**

**MTP**

**LTP**

**Supplementary Figure 5) 12 months old *Mig-6^over/over^* female mice showed little damage.** Representative images of Toluidine Blue stained sections of knee joints from 12-month female control **(A)** and female Mig-6 over **(B)** mice were evaluated for cartilage damage following OARSI histopathological scale on the four quadrants of the knee: LFC = lateral femoral condyle, LTP = lateral tibial plateau, MFC = medial femoral condyle and MTP = medial tibial plateau. OARSI based cartilage degeneration scores are higher both in the MFC and MTP of Mig-6 overexpressing mice, corresponding to the increased damage observed histologically **(C)**. Data analyzed by two-way ANOVA with Bonferroni's multiple comparisons test. Individual data points presented with mean ± SEM. All scale bars =100 μm. N = 8 mice/group. Scale bar = 100µm.

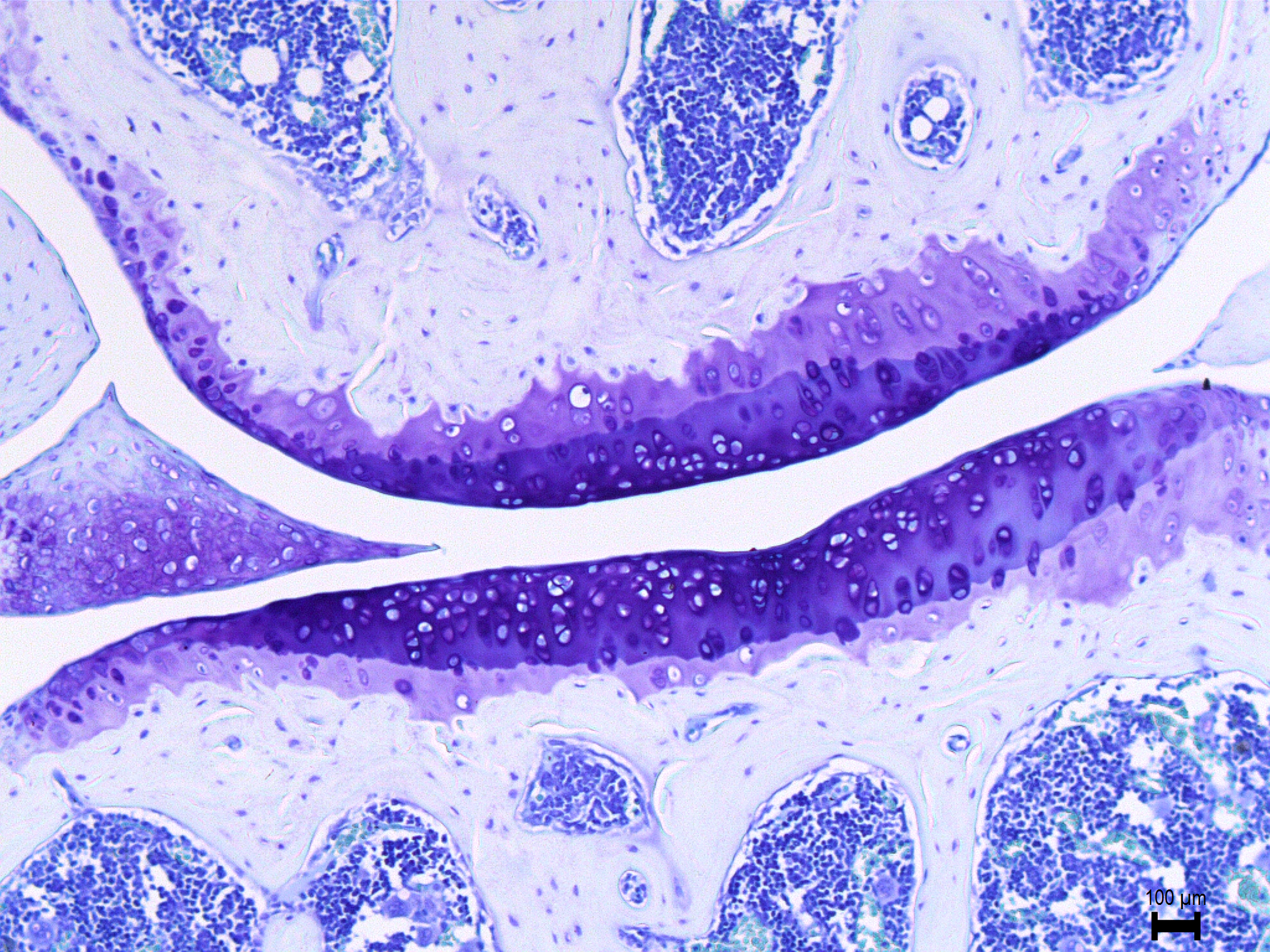

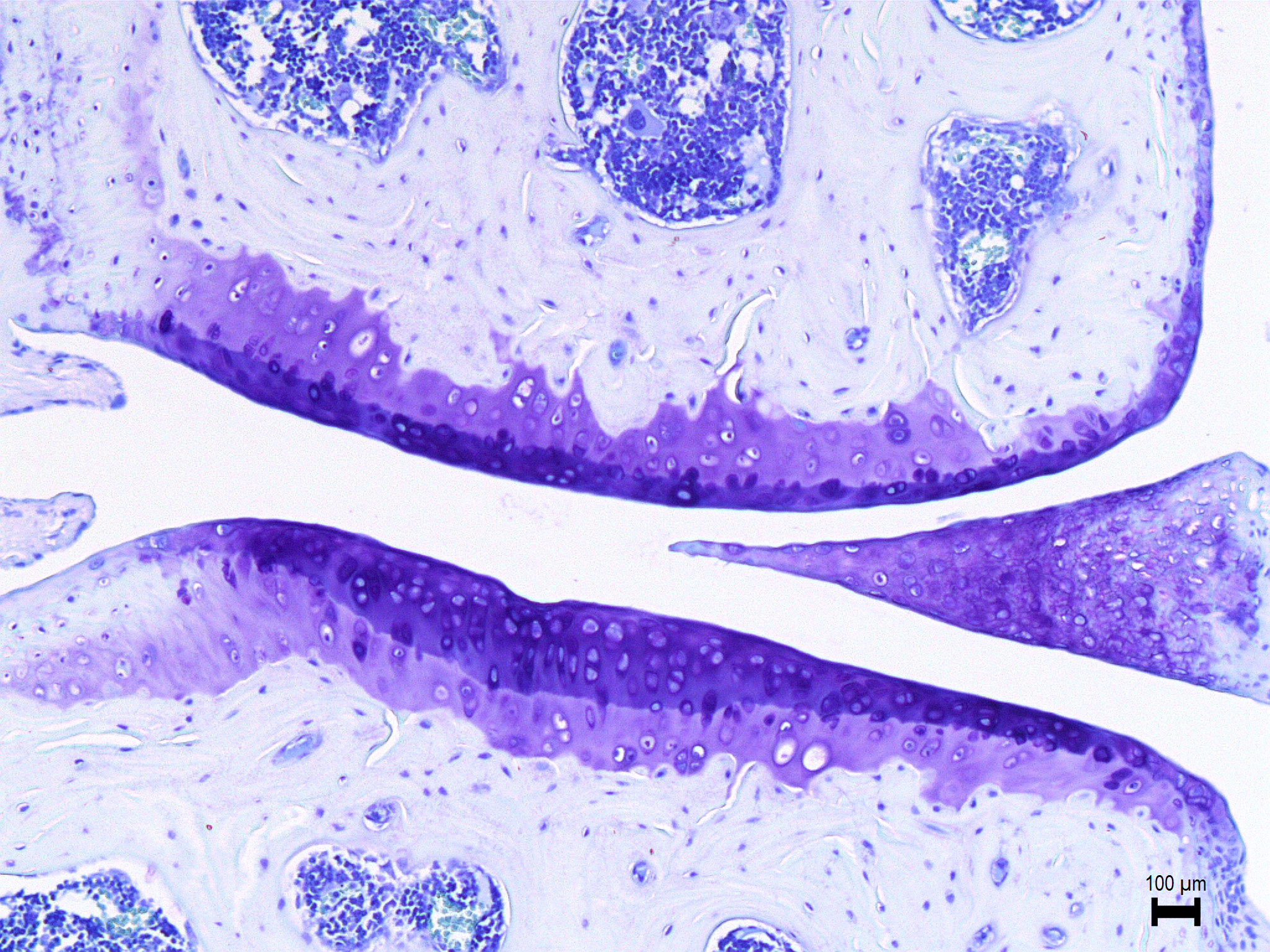

A

**Control**

**18 months old**

**MTP**

**LTP**

**LFC**

**MFC**

C

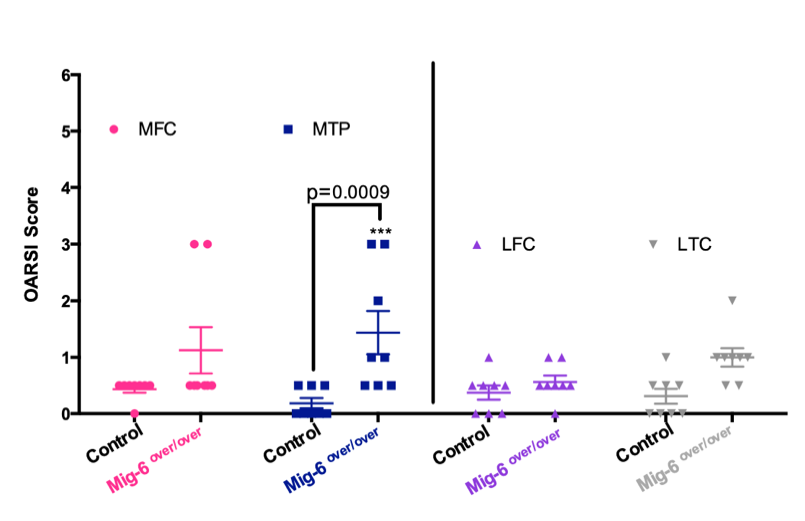

***Mig-6^over/over^***

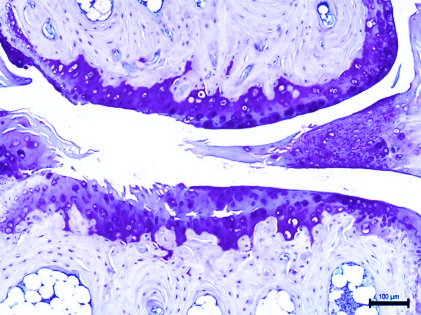

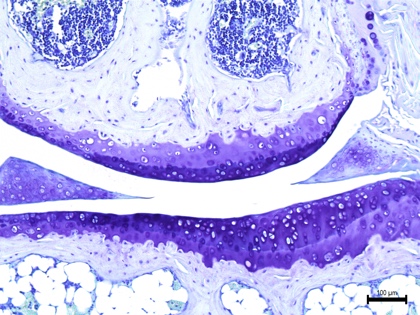

B

**MFC**

**MTP**

**LFC**

**LTP**

**Supplementary Figure 6) 18 months old *Mig-6^over/over^* female mice showed little damage.** Representative images of Toluidine Blue stained sections of knee joints from 18-month female control **(A)** and female Mig-6 over **(B)** mice were evaluated for cartilage damage following OARSI histopathological scale on the four quadrants of the knee: LFC = lateral femoral condyle, LTP = lateral tibial plateau, MFC = medial femoral condyle and MTP = medial tibial plateau. OARSI based cartilage degeneration scores are higher both in the MFC and MTP of Mig-6 overexpressing mice, corresponding to the increased damage observed histologically. **(C)**. Data analyzed by two-way ANOVA with Bonferroni's multiple comparisons test. Individual data points presented with mean ± SEM. All scale bars =100 μm. N = 8 mice/group. Scale bar = 100µm.

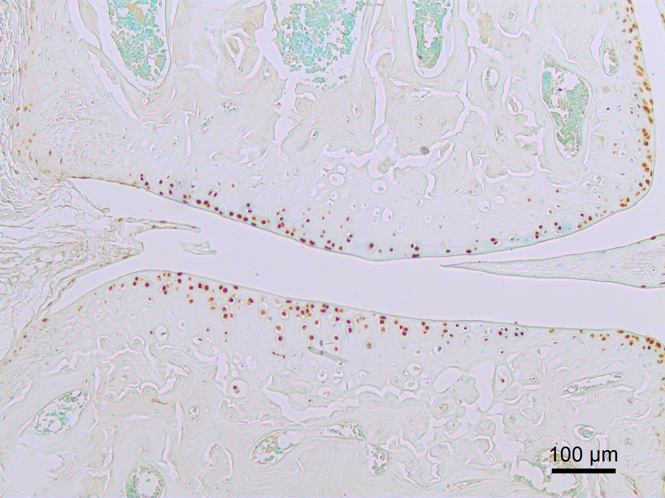

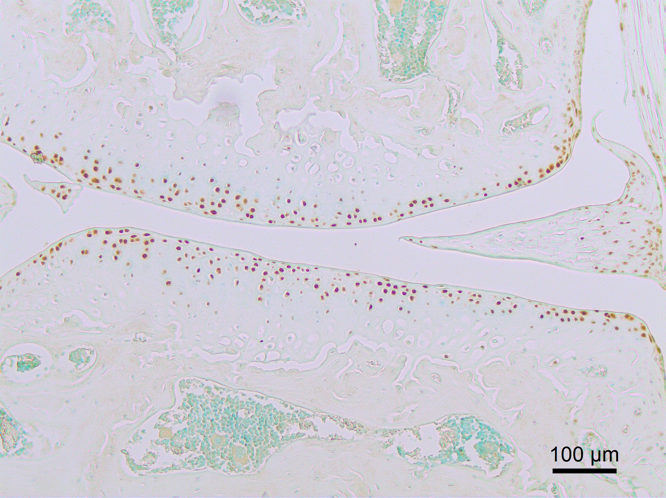

***Mig-6^over/over^***

**Control**

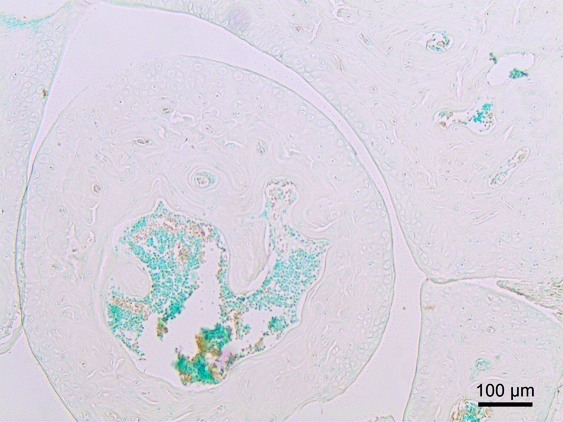

**No primary antibody control**

A

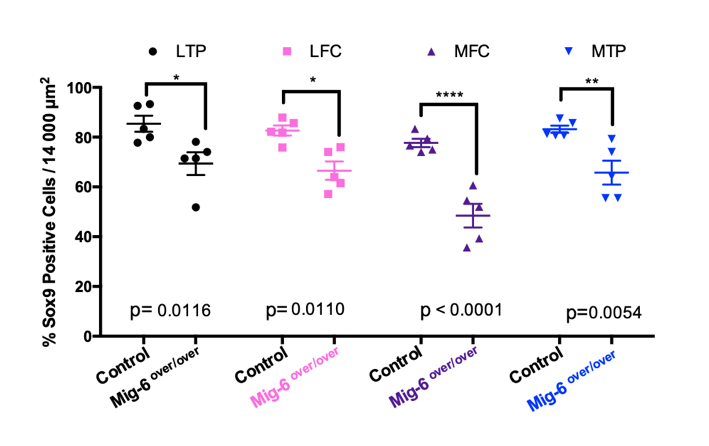

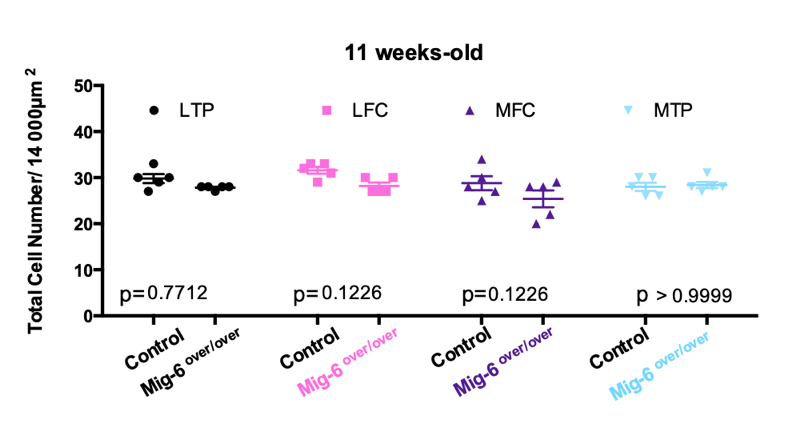

B

**Supplementary Figure 7) SOX9 immunostaining shows a decrease in Mig-6 overexpressors mice at 11 weeks-old male mice control and Mig-6over. No primary antibody staining.** Ratio between the total cell number from control and Mig-6over at 11 weeks-old male mice **(A).** Ratio between the percentage of Sox9 positive cells from control and Mig-6over at 11 weeks-old male mice **(B).** Data analyzed by two-way ANOVA (95% CI) with Bonferroni post-hoc test. Individual data points presented with mean ± SEM; N= 5 mice/genotyping. LFC = lateral femoral condyle, LTP = lateral tibial plateau, MFC = medial femoral condyle and MTP = medial tibial plateau. Scale bar = 100µm.
